# Connectome analysis of a cerebellum-like circuit for sensory prediction

**DOI:** 10.1101/2025.07.03.662989

**Authors:** Krista E. Perks, Mariela D. Petkova, Salomon Z. Muller, Michael Genecin, Adishree Ghatare, Richard Schalek, Yuelong Wu, Michal Januszewski, Viren Jain, Jeff W. Lichtman, LF Abbott, Nathaniel B. Sawtell

## Abstract

Stable and accurate perception involves comparing incoming sensory input with internally-generated predictions ^1–3^. A mechanistic understanding of this process has been elusive due to the size and complexity of the relevant brain regions in mammals. Here we leverage connectomics to comprehensively map the cell types and synaptic connections underlying a well-characterized and ecologically relevant form of predictive sensory processing in the cerebellum-like electrosensory lobe (ELL) of weakly electric fish ^4,5^. Connectome analysis reveals highly-structured feedforward and recurrent synaptic connectivity mediating the cancellation of predictable electrosensory input. A computational model constrained by prior electrophysiological recordings shows how this connectivity supports the formation of predictions at multiple sites within the network and how the ELL solves a continual learning problem by maintaining fast and accurate predictions despite noise and changes in environmental context. Overall, these findings provide a blueprint for using connectomics to elucidate learning in vertebrate nervous systems.

## Main

Comparing incoming sensory input with internally-generated predictions allows organisms to distinguish behaviorally relevant stimuli from those that are self-generated ^6–9^. One particularly well-characterized example is a species of African freshwater fish that both senses and emits electrical fields. The emitted fields, brief pulses known as electric organ discharges (EOD), are used for communication and active sensing. However, EODs interfere with a separate passive electrosensory system that uses a class of low-frequency tuned electroreceptors on the fish’s skin to sense weak electrical fields emitted by the fish’s prey ^10^. This interference problem is solved by a cerebellum-like structure in the hindbrain of the fish known as the electrosensory lobe (ELL) ^5^. The ventrolateral zone of the ELL (hereafter referred to simply as the ELL) integrates sensory input from passive electroreceptors with copies of the command to discharge the electric organ ^4,11^ (Figure 1A,B) . These copies elaborate the command signal into a temporally diverse and delayed set of granule cell inputs that are sculpted by synaptic plasticity into a temporally-specific prediction of the sensory impact of the self-generated EOD ^12–16^. This prediction is subtracted from the sensory input stream, allowing the fish to detect weak external fields due to prey ^17^. Synaptic plasticity is vital for maintaining the accuracy of this prediction in the face of changing environmental factors ^10^ and noise.

**Figure 1.**
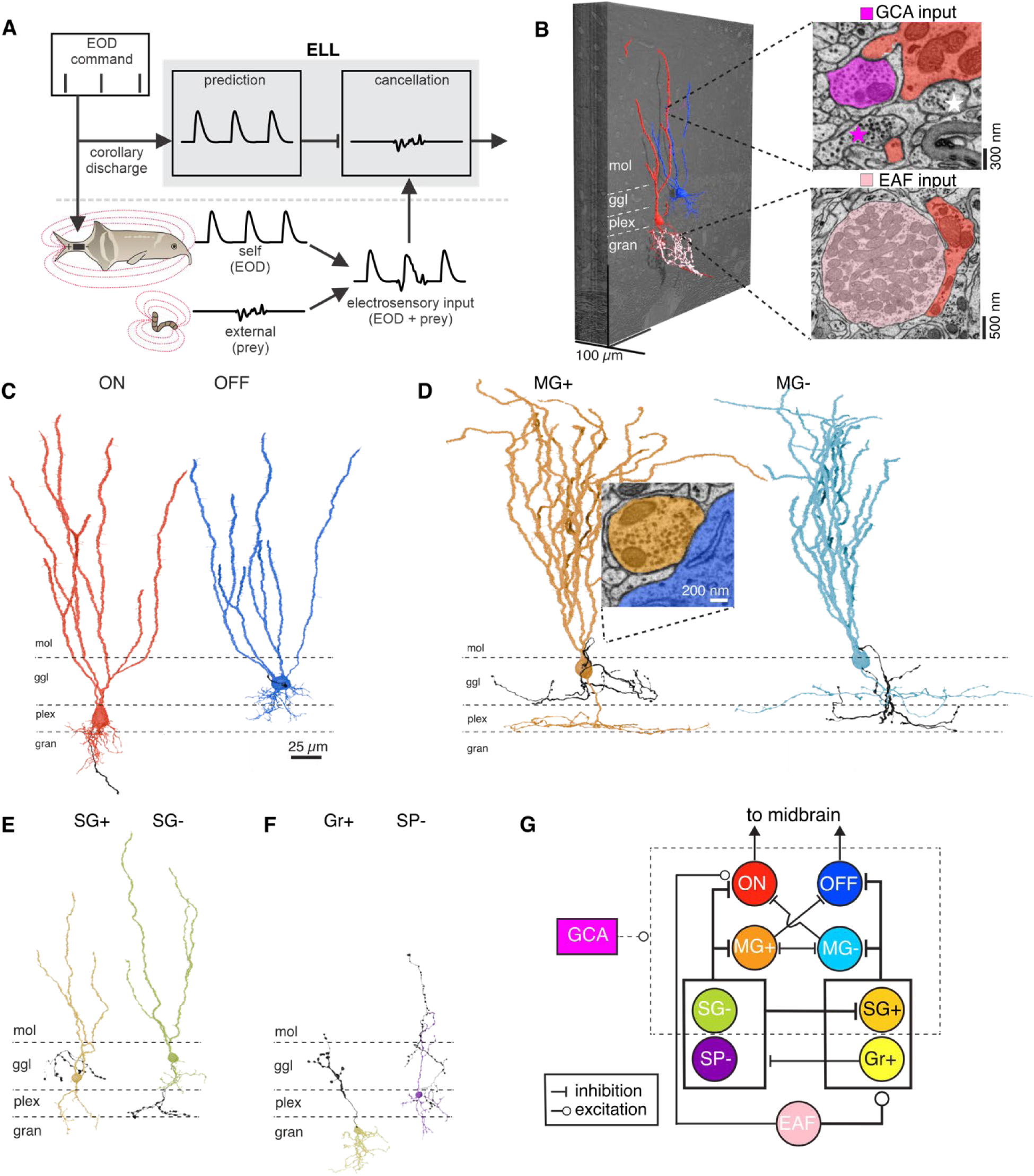
Connectome-based analysis of cell types and circuitry of the ELL. **A**, The ELL integrates input from electroreceptors with a diverse array of signals conveyed by granule cells, including electric organ corollary discharge signals. Electrophysiological studies have shown that temporally-specific predictions within the ELL cancel self-generated components of the electrosensory input, revealing behaviorally- relevant signals. **B**, Cutout (412 x 409 x 48 µm) from the full imaged EM volume (450 × 450 × 105 µm) showing 2 reconstructed cells (blue and red) and EAF (pink) and GCA (magenta) inputs. The volume contains the central sensory representation of a small portion of the anterior body surface (see Methods). The dorso-ventral and anterior-posterior dimensions of the skin surface are mapped onto the *x* and *z* dimensions of the volume, respectively. Top inset: excitatory synapses (magenta) onto apical dendritic spines of an Output ON cell (red fill). Magenta fill shows the reconstructed GCA visualized in the volume; magenta star marks another GCA in the EM section. White star marks an inhibitory molecular layer interneuron synapse onto the dendritic shaft of the same cell. Bottom inset: Excitatory EAF synapse (pink fill) onto the basal dendrite of an Output ON cell (red fill). mol, molecular layer; ggl, ganglion layer; plex, plexiform layer; gran, granular layer. **C-F**, Reconstructions of major ELL cell types. ON: Output ON, OFF: Output OFF, MG+/MG-: medium ganglion, SG+/SG-: small ganglion, Gr+: granular, SP-: small plexiform. Inset: example inhibitory MG+ synapse onto an Output OFF cell. **G**, Summary diagram of the synaptic connectivity patterns revealed in the present study. Only connections comprising >10% of the total synapses to a given cell type are shown in the diagrams. Inputs and outputs shown to/from boxes around multiple cell types indicate common connections among those cell types.

Here we leverage connectomics, in tight combination with computational modeling, to provide detailed answers to two questions of broad relevance for predictive processing ^9^. First, where and how are predictions generated and, second, where and how are predictions compared with sensory input? Extensive prior *in vitro* and *in vivo* electrophysiological recordings and modeling suggest a particular implementation of these functions by specific cell types within the ELL ^12–16,18,19^. This hypothesized implementation depends critically on specific patterns of synaptic connectivity that can be directly tested using connectomics. Predictive sensory processing in the ELL is uniquely amenable to connectomic analysis because the major elements involved in the computation are identifiable solely on the basis of their morphology, they have been subject to extensive prior electrophysiological and computational analysis, and they are restricted to a small tissue volume ^20–27^ (although some synapses in our study cannot be characterized - see Discussion).

Our analysis centers on two major cell types of the ELL that have been the focus of prior electrophysiological recording and modeling studies: glutamatergic projection neurons, termed Output cells, and Purkinje-like interneurons, known as medium ganglion (MG) cells, that inhibit Output cells ^21^ (Figures 1C,D). Each of these types is sub-divided into two morphologically and electrophysiologically distinct sub-types (Output ON, Output OFF, MG+, and MG-). Output ON and MG+ cells respond to electrosensory stimuli with the same polarity as electroreceptors, whereas Output OFF and MG- respond with the opposite polarity ^19,28^. We discuss the operation of the ELL in a modeling section below, but, in brief, MG cells extract, through learning, a prediction of the sensory consequences of the EOD. MG inhibition, along with additional learning that takes place within the Output cells, results in a subtraction of the predictable, EOD-related signal from the sensory input stream (Figure 1A).

Before diving into our connectome study, we present a brief summary of our major findings to help guide the reader through the level of detail involved in our analysis. This level of detail is, in fact, the point of our study: we provide here a view of a circuit for predictive learning "even unto its innermost parts". The major results from this analysis are:

1. Input pathways to ON and OFF sub-circuits are highly segregated (Figure 1G).
2. Although the sensory afferents are excitatory, MG+ cells receive most of their sensory input through a disynaptic inhibitory pathway rather than directly. MG- cells receive sensory input through a single inhibitory stage (Figure 1G).
3. The connectivity of MG cells is precisely structured, with MG+ cells inhibiting Output OFF and MG- cells, and MG- cells inhibiting Output ON and MG+ cells.
4. We use our connectome results and previous neural recording data to construct a highly constrained model of learning in the ELL. This model explains the logic of the major connectome features listed in 1-3 above, shows that the ELL learns at two different speeds, and illustrates the advantages of this two-speed learning system under naturalistic conditions.

### Serial-section EM reconstruction of ELL circuitry

We generated a 450 × 450 × 105 µm EM volume from an anterior portion of the ELL using serial section electron microscopy with the ATUM-mSEM approach developed for a human cortex dataset ^29^ (Figures 1B and Extended Data Figure 1). Automated segmentation was performed using a flood filling network (FFN) pipeline ^30^ that was retrained for the ELL volume, followed by manual synapse annotation (see Methods). Consistent with prior EM studies of the mormyrid brain, ultrastructure and synapse morphology resembled that of other vertebrates, including typical ‘asymmetric’ (putative excitatory) and ‘symmetric’ (putative inhibitory) synapses ^31,32^ (Figures 1B,D and Extended Data Figure 1).

The ELL integrates two main streams of excitatory input, both of which could be clearly identified within the volume: granule cells axons (GCAs) and electroreceptor afferent nerve fibers (EAFs). GCAs travel long distances in the molecular layer of the ELL (similar histologically to the cerebellar molecular layer ^33^) synapsing onto the spiny apical dendrites of MG and Output cells (Figure 1B). The cell bodies of granule cells are located in a cell mass, the eminentia granularis posterior (EGp), located outside of the reconstructed volume (Extended Data Figure 1). EAFs innervate electroreceptors on the skin and project somatotopically to the ELL, synapsing onto interneurons in the granular layer (Figure 1B). The electrophysiological response properties of both EGp granule cells and EAFs have been characterized extensively in prior studies, making the inputs to the ELL well understood ^10,15,28,34^.

Since our main goal was to test specific hypotheses, we focused our efforts on targeted reconstructions of the synaptic inputs and outputs for specific cell types of interest rather than attempting to densely reconstruct the entire volume (which contains on the order of 10,000 neurons). To initiate this process, we started by reconstructing large-soma cells in the ganglion layer known from previous work to be MG and Output cells, and then worked backward and forward from synaptic annotation of these key cell types. The Output ON, Output OFF, MG+, and MG- cells mentioned above could all be classified unambiguously using a combination of manual sorting and automated clustering based on quantitative measurements of various morphological features (Extended Data Figure 2). Iterative annotation and reconstruction of MG cell synaptic targets resulted in a densely reconstructed network of 88 Output and 136 MG cells. Tracing the input pathways to these two main cell types resulted in the sparse reconstruction of three additional interneuron types: small ganglion (SG), granular (Gr+), and small plexiform (SP-) cells (Figures 1E,F and Extended Data Figure 2). Like the MG cells, SG cells are subdivided into subclasses referred to as SG+ and SG- on the basis of their axonal and dendritic morphologies (Extended Data Figure 2). Although the *in vivo* response properties of SG, GR+, and SP- cells have not been characterized, their synapse morphology along with prior immunohistochemical and EM studies indicate that they are inhibitory ^20,21,23,25^ (Extended Data Figure 1). In total, our analysis includes 17,348 annotated synapses and 1,749 reconstructed and classified neurons.

### The granule cell input pathway

Granule cells of the EGp convey EOD command signals (as well as additional information) that provide the basis for predicting self-generated sensory input to the ELL^15,34^. To estimate the relative proportion of GCA input received by different ELL cell types, we densely reconstructed all of the neurons with spiny apical dendrites (Output, MG, and SG cells) located within a small subvolume of the ganglion and plexiform layers (60 x 60 x 60 microns) and performed skeletonization to quantify their apical dendritic lengths in the molecular layer (Figure 2A).

**Figure 2.**
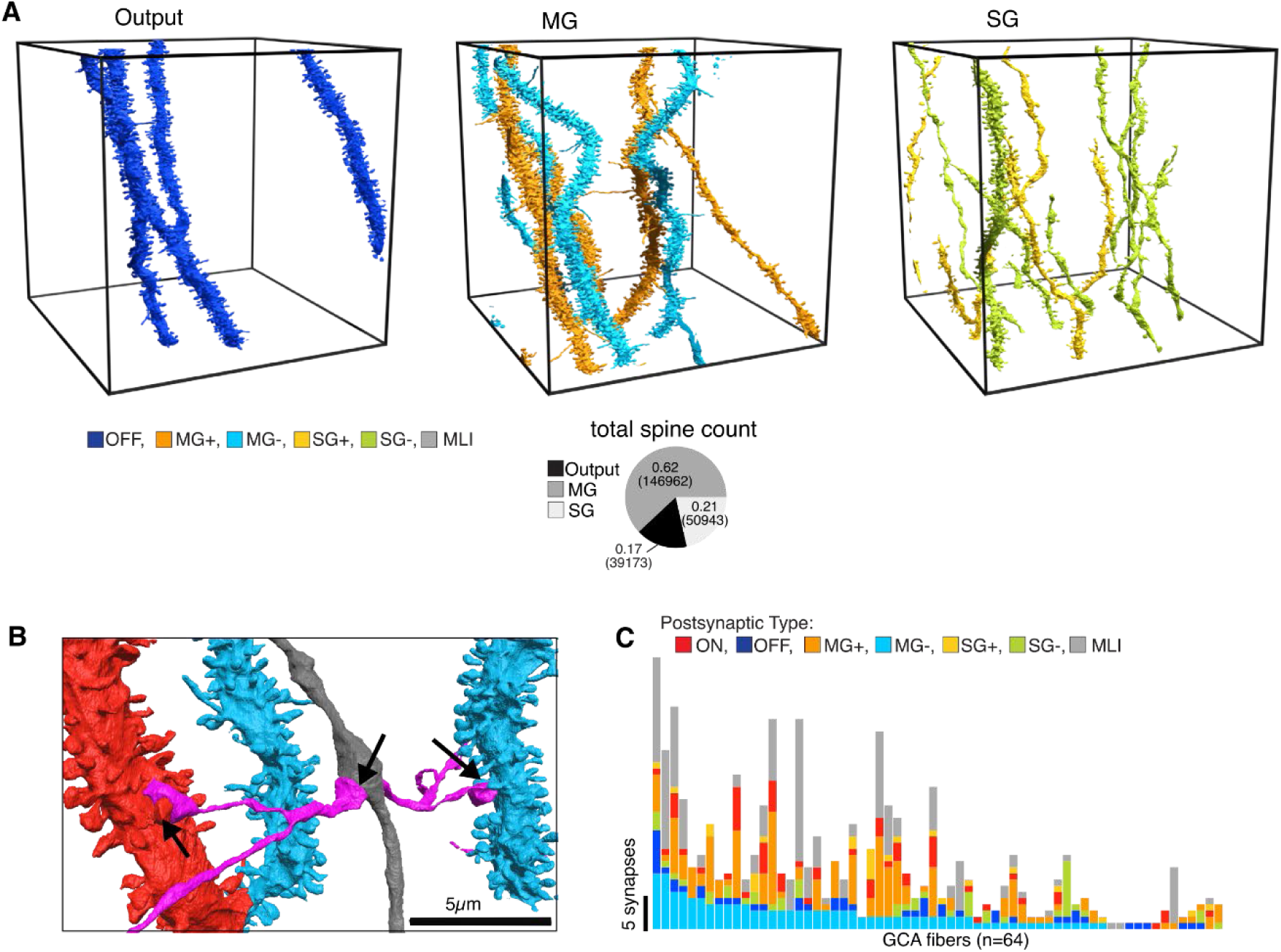
The granule cell input pathway. **A**, Apical dendrites in the molecular layer (∼100 µm from the surface of the ganglion layer) corresponding to all of the Output, MG, and SG cells located in a 60 x 60 x 60 µm subregion of the EM volume. Pie chart shows the estimated fraction of spines per cell type (and estimated total spine count) corresponding to the 6 Output, 13 MG, and 93 SG cells within the selected subvolume. **B**, A GCA (magenta) synapsing onto the spines of an Output ON (red) and MG- (blue) cell and onto the swelling of a smooth apical dendrite (gray) (arrows show synapse locations). **C**, Number of synapses formed by each reconstructed and synapse-annotated GCA onto different postsynaptic cell types (indicated by colors) (n = 676 synapses from 64 GCA onto 312 Output, MG, SG, and MLI neurons). GCA fibers are ordered along the x-axis by the number of synapses onto MG- (decreasing) and then MG+ (increasing). An additional 321 GCA synapses were onto unclassified elements.

Combining the dendritic length measurements with separately obtained spine density estimates, yielded the relative proportions of total GCA input onto all of the reconstructed cells of a particular type: collectively the MG cells receive 3.6 times and the SG cells 1.2 times as much total GCA input as the Output cells (Figure 2A). By comparison, estimated median spine counts for individual MG cells were 14,000, compared to 7,000 for Output and 350 for SG cells.

We next analyzed the connectivity of GCAs (n = 64), which were identified as extremely thin, unmyelinated fibers that traveled for long distances in the molecular layer, often exiting the volume. GCAs form synapses onto dendritic spines of Output, MG, and SG cells as well as onto the smooth, beaded dendrites of molecular layer interneurons (MLIs; Figures 2B). Individual GCAs synapse onto various combinations of Output, MG, and SG cells of both sub-types (Figure 2C). Although we cannot rule out subtle biases in GCA connectivity, these results are consistent with the view that unstructured input from granule cells provides a rich basis set of predictive signals that are subsequently sculpted by activity-dependent synaptic plasticity. The prominence of SG cells was unexpected and suggests a previously unappreciated role for these cells in predictive processing (see Discussion).

### Electrosensory input pathways

Our next goal was to map the electrosensory input pathways through the ELL. Specifically, we sought to reveal the anatomical basis for the separation of Output and MG cells into ON and OFF subtypes based previously on their physiological responses to electrosensory stimuli. *In vivo* recordings have shown that EAF input to MG cells strongly modulates dendritic spiking while minimally impacting axonal spike rates^18^. These selective effects of sensory input are vital for allowing MG cells to simultaneously generate predictions (via plasticity induced by dendritic spiking) and transmit them to Output cells (via axonal spike rate modulations). Prior biophysical modeling suggests these functional requirements can be met if electrosensory input to MG cells is relayed by inhibition and dis-inhibition ^18,19^. Because EAFs are excitatory, the anatomical basis for such specialized sensory pathways was unclear.

Arrows show the locations of the 4 cells used for targeted reconstructions of presynaptic partners shown in **C** and **D**. **C**, Left: locations of example Output ON and OFF cells used for targeted reconstructions of presynaptic partners. Middle: all categorized presynaptic partners (excluding MG) to the Output ON cell (n = 240 total synapses from SG-, SP-, and EAF neurons; 574 total synapses were annotated, with 15% of those synapses from MG cells that will be analyzed separately and 43% unclassified). Right: all categorized presynaptic partners (excluding MG) to the Output OFF cell (n = 363 total synapses from SG+ and Gr+ neurons; 530 total synapses were annotated, with 8% of those synapses from MG cells that will be analyzed separately and 23% unclassified). **D**, Left: locations of the MG+ and MG- cells used for targeted reconstructions of presynaptic partners. Middle: all categorized presynaptic partners (excluding MG) to the MG+ cell (n = 238 total synapses from SG-, SP-, and EAF neurons; 460 total synapses were annotated, with 11% of those synapses from MG cells that will be analyzed separately and 37% unclassified). Right: all categorized presynaptic partners (excluding MG) to the MG- cell (n = 217 total synapses from SG+ and Gr+ neurons; 355 total synapses were annotated, with 10% of those synapses from MG cells that will be analyzed separately and 29% unclassified). **E**, Number of synapses from each cell type onto individual Output and MG cells (n = 2838 total synapses from 925 SG, SP-, Gr+, and EAF neurons; 5003 total synapses were annotated, with 10% of those synapses from MG cells that will be analyzed separately and 33% unclassified). Cells are ordered along the horizontal axis by the number of inputs from Gr+ (decreasing) and then SP- (increasing). Complete reconstructions of all presynaptic partners were made for one cell of each type. Reconstruction of a random subset of presynaptic partners were performed for additional example cells of each type. Pie charts show the fraction (and total number) of synapses from each cell type onto all ON, OFF, MG+ and MG- cells represented in the bar graphs. **F**, Observed connectivity from EAF, Gr+, SP-, SG+, and SG- to Output cells compared with that expected from random connectivity applied to the 10 output cells (5 ON, 5 OFF) shown in **E**. Each entry corresponds to input from the cell type shown on the horizontal axis given that the postsynaptic cell receives at least one synapse from the cell type shown on the vertical axis (see Methods). Presynaptic cell types are grouped along these axes according to their responses inferred from connectivity and/or known from previous *in vivo* recordings. Two clusters (illustrated with dotted lines) are observed among the input cell types, corresponding to the ON and OFF pathways. **G**, same as in **F** for the 10 MG cells (5 MG-, 5 MG+) shown in **E**.

To trace electrosensory input pathways, we chose example Output ON, Output OFF, MG+, and MG- cells and reconstructed the cells synapsing onto their basal dendrites, soma, axon, and proximal apical dendrites (Figure 3A,B). This analysis excludes synapses onto apical dendrites, as these mainly come from GCAs, and synapses from MG cells, which are discussed separately below. Note that the different cell types of the ELL, including those chosen here for reconstruction of presynaptic inputs, are intermixed in the x and z dimensions of the volume without any obvious spatial pattern (Figure 3B).

**Figure 3.**
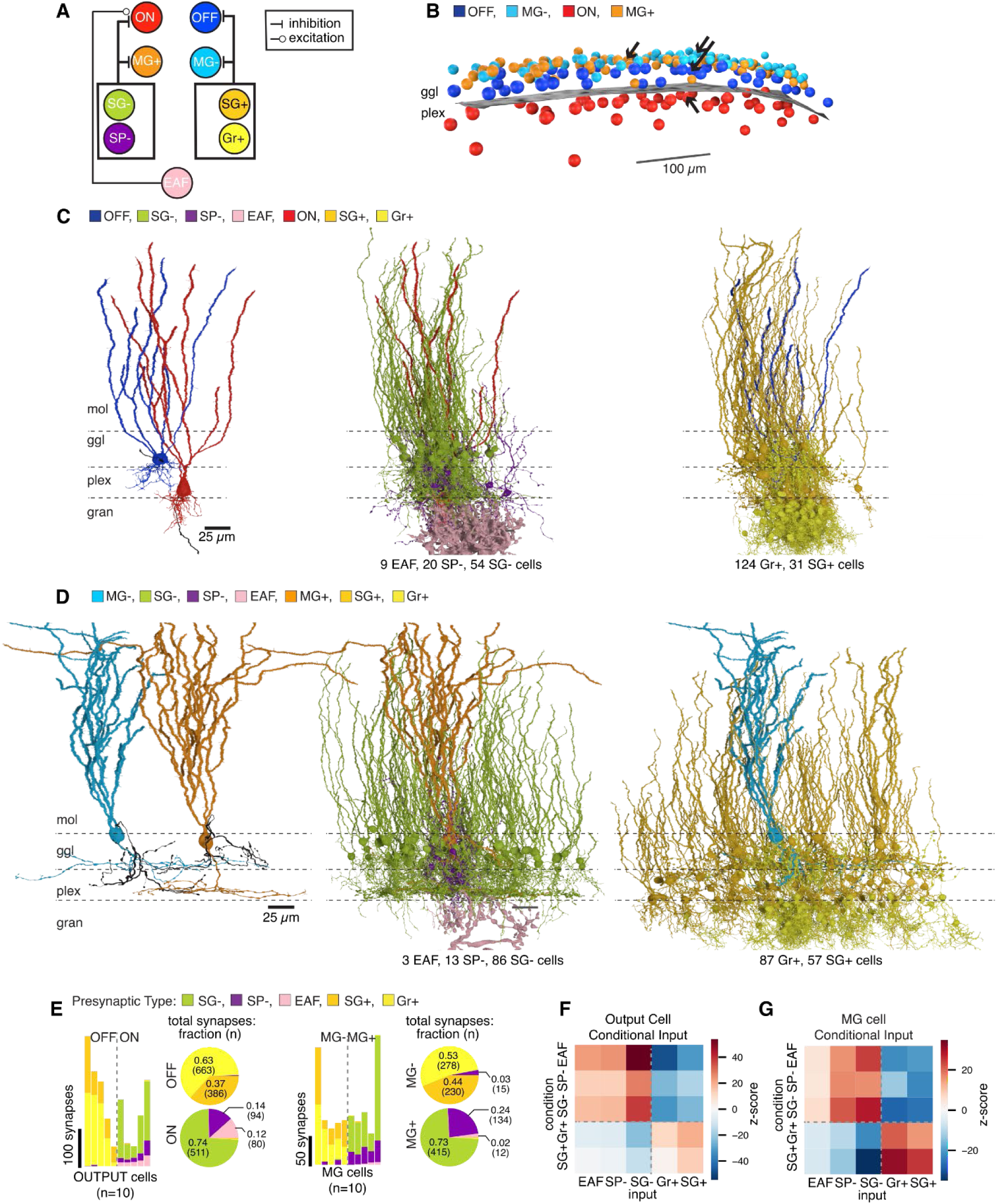
Sensory input pathways to Output and MG cells. **A**, Main cell types synapsing onto Output and MG cells (excluding MG cells and synapses formed in the molecular layer). **B**, Visualization of the soma locations of all reconstructed Output (42 OFF and 46 ON) and MG (67 MG- and 69 MG+) cells in the EM volume. Plane shows the boundary between the plexiform (plex) and ganglion (ggl) layers.

Putative excitatory synapses from EAFs made up only a small fraction of the input to Output ON and MG+ cells. The overwhelming majority of input to all 4 cell types arose instead from the SG+, SG-, Gr+, and SP- inhibitory interneuron classes (Figure 3C,D). A strictly segregated pattern of connectivity was observed in which Output ON and MG+ cells received synapses from SG-, and SP-, and EAFs, whereas Output OFF and MG- cells received synapses from SG+ and Gr+ cells (Figures 3E-G and Extended Data Figure 3).

Proceeding backwards along the electrosensory pathway, we reconstructed presynaptic neurons synapsing onto SG+, SG-, SP-, and Gr+ cells (Figure 4A). Again, a strictly segregated pattern of connectivity was observed in which SG- and SP- cells received the large majority of their input from Gr+ cells, whereas SG+ cells received both EAF and SG- input (Figure 4B-F). Finally, Gr+ cells received the large majority of their input from EAFs (Figure B,D,F).

**Figure 4.**
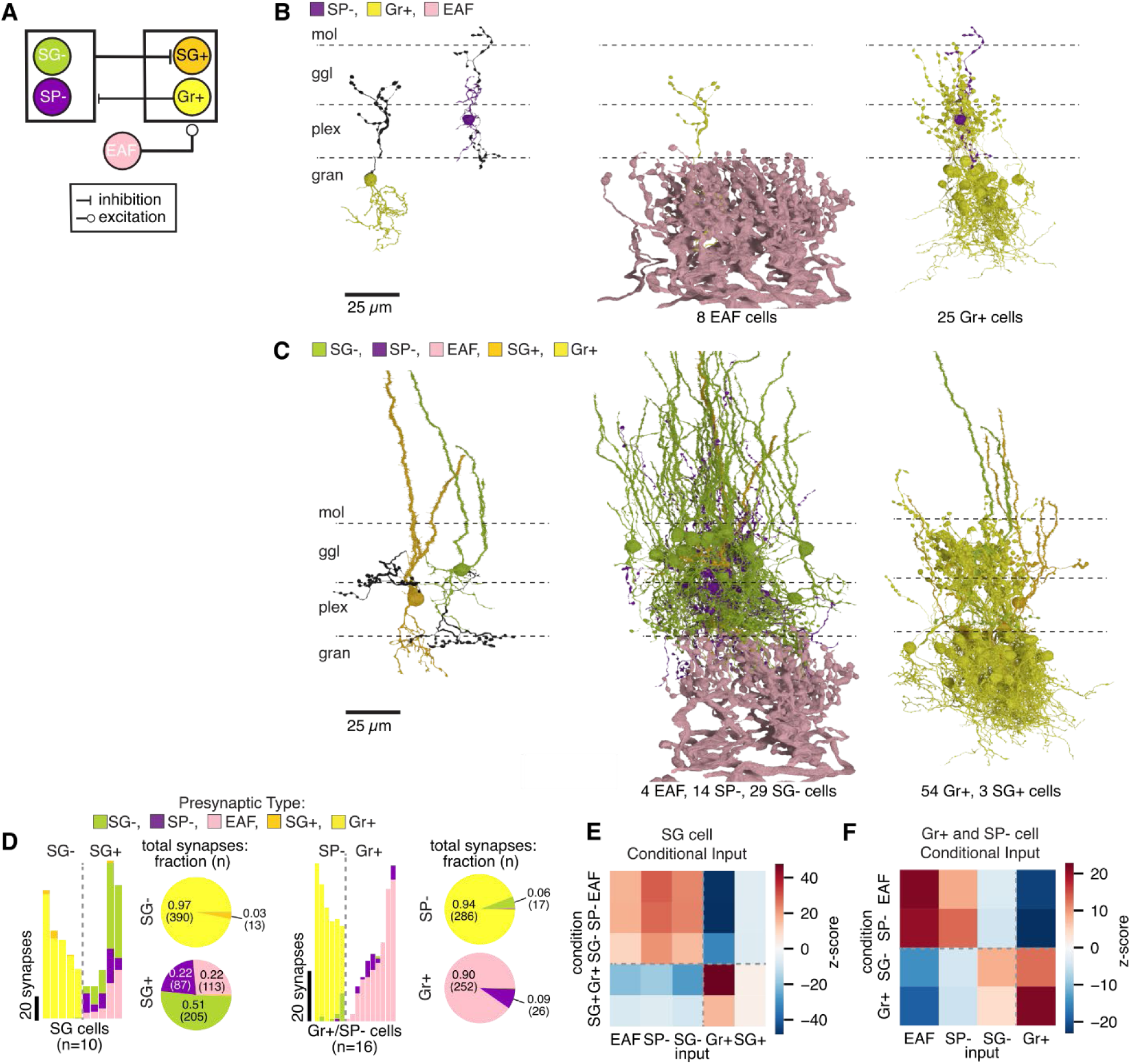
Sensory input is conveyed by inhibitory and dis-inhibitory pathways. **A**, Summary diagram of the main cell types synapsing onto SG, Gr+, and SP- cells. **B**, Left: locations of the Gr+ and Sp- cells used for targeted reconstructions of presynaptic partners. Middle: all categorized presynaptic partners to the Gr+ cell (n = 61 total synapses from EAF neurons; 216 total synapses were annotated, with 3% of those synapses from SP cells and 69% unclassified). Right: all categorized presynaptic partners to the SP- cell (n = 42 total synapses from Gr+ neurons; 55 total synapses were annotated, with 1 synapse from an SG cell and 22% unclassified). **C**, Left: locations of the SG+ and SG- used for targeted reconstructions of presynaptic partners. Middle: all categorized presynaptic partners to the SG+ cell (n = 156 total synapses from SG-, SP-, and EAF neurons; 201 total synapses were annotated with 22% unclassified). Right: all categorized presynaptic partners to the SG- cell (n = 127 total synapses from SG+ and Gr+ neurons; 157 total synapses were annotated, with 20% unclassified). **D**, Number of synapses from each cell type onto individual SG, Gr+ and SP- cells (n = 1387 total synapses from 457 SG, SP-, Gr+, and EAF neurons; 2366 total synapses were annotated, with 41% unclassified). Cells are ordered along the horizontal axis by the number of inputs from Gr+ (decreasing) and then EAF (increasing). Pie charts show the fraction and total number of synapses from each cell type onto all SG+, SG-, Gr+, and SP- cells represented in the bar graphs. **E**, Observed connectivity from EAF, Gr+, SP-, SG+, and SG- to SG cells compared to that expected based on random connectivity. Conditional input analysis for the 11 SG cells (5 SG-, 6 SG+) represented in **D**. Each entry corresponds to input from the cell type shown on the horizontal axis given that the postsynaptic cell receives at least one synapse from the cell type shown on the vertical axis (see Methods). Presynaptic cell types are grouped along these axes according to their responses inferred from connectivity and/or known from previous *in vivo* recordings. Two clusters (illustrated with dotted lines) are observed among the input cell types, corresponding to the ON and OFF pathways. **F**, Same display as in **E** for the 6 SP- and 10 Gr+ cells represented in **D**.

Summarizing these connectivity patterns, two serial stages of inhibition (along with some direct EAF excitation) preserve the sign of EAF input to Output On and MG+ cells, whereas a single stage of inhibition inverts the sign of EAF input to Output OFF and MG- cells (Figure 4A). These results provide an anatomical basis for the distinct ON and OFF responses to sensory input observed in Output and MG cell sub-types (major result 1 in the Introduction) as well as striking support for the hypothesized dis-inhibitory and inhibitory effects of electrosensory input on MG cells (major result 2 in the Introduction).

### Synaptic connectivity underlying the transmission of sensory predictions

Prior *in vivo* intracellular recordings and modeling suggests that the cancellation of self-generated sensory input in Output cells is mediated, in part, by predictions transmitted by MG cells ^18^. A specific wiring scheme is required to achieve this. Because of the way predictive signals are learned (see modeling section below), the predictive signal appropriate for an Output ON cell is generated by MG- cells, and the appropriate predictive signal for an Output OFF cell is generated by MG+ cells (Figure 5A). To test whether this requirement is satisfied in the connectome, we annotated all of the MG cell synapses in the volume (n = 7,661) and reconstructed their postsynaptic partners (n = 1,155). MG cells overwhelmingly targeted Output cells (∼71 % of synapses onto classified targets) and other MG cells (∼27 % of synapses classified targets) (Figure 5B). Moreover, their synaptic connectivity (which was annotated blind to cell type identity) obeyed a remarkably strict pattern: MG- cells synapsed onto Output ON cells and MG+ cells, whereas MG+ cells targeted Output OFF cells and MG+ cells (major result 3 of the Introduction; Figure 5B-E and Extended Data Figure 4). The accuracy of this synaptic targeting was remarkable: of a total of 3,989 synapses from MGs to Output cells, only 1 synapse violated the - to ON, + to OFF pattern. Similarly, of 1521 MG-to-MG synapses only 11 were between similar MG subtypes. This is a testament both to the accuracy of synaptic targeting in the ELL and to the quality of our connectome reconstructions. Although the spatial overlap between MG cell axons and Output cell somata suggest a rough wiring selectivity ^18^, it cannot account for the nearly perfect wiring specificity demonstrated here. Additional connectome analysis revealed that MG synapses are not restricted to Output cell somata, but also synapse heavily on their basal dendrites (Figure 5H). Similarly, the highly-selective recurrent connections between MG sub-types we discovered could not be inferred on the basis of the soma-axon overlap, because both MG+ and MG- cell somata are located in the ganglion layer (Figure 5I-J). The functional significance of this recurrent connectivity between MG cells will be analyzed further below.

**Figure 5.**
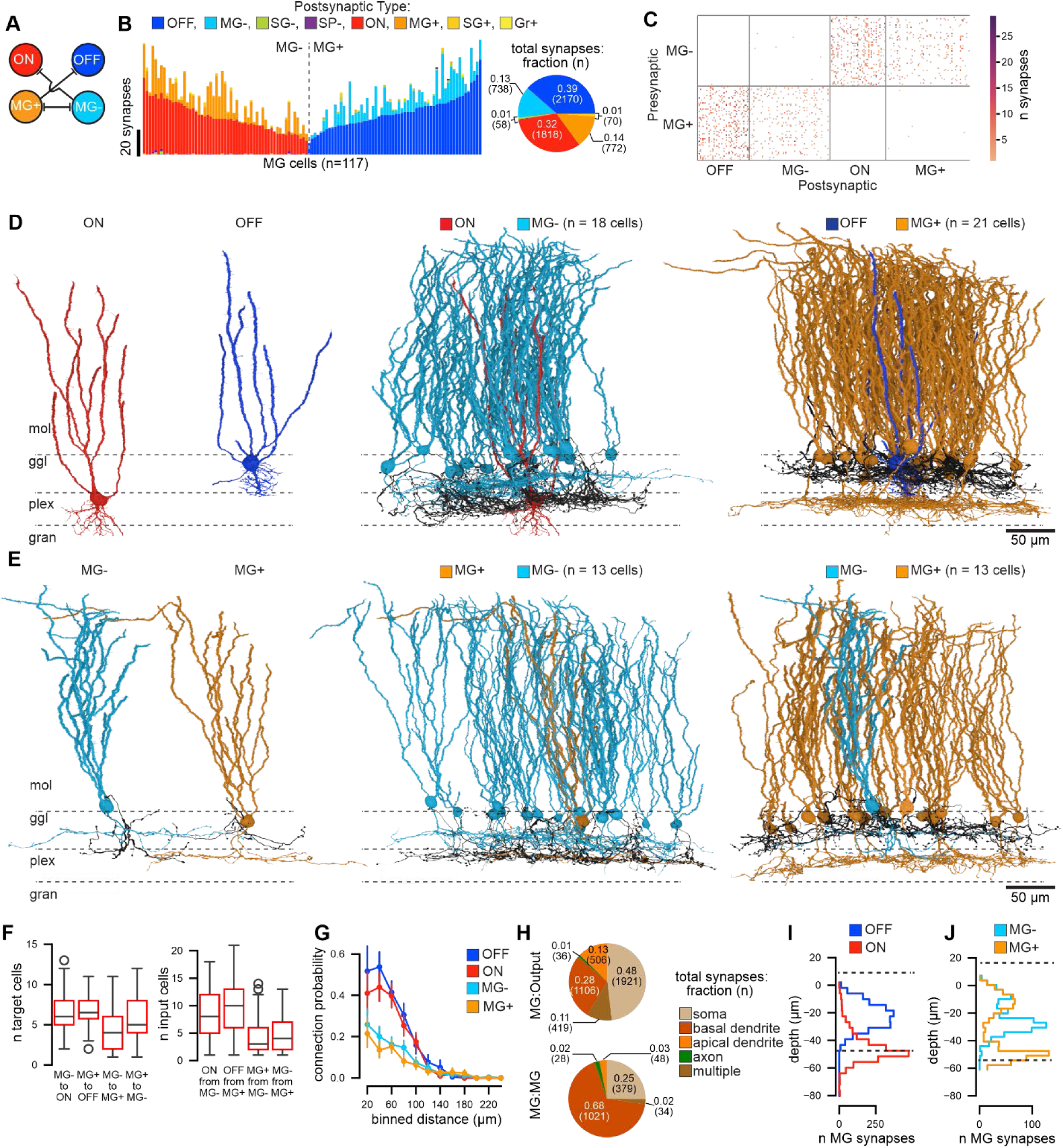
Feedforward and recurrent connectivity of MG cells. **A**, Feedforward and recurrent connectivity between MG and Output cell subtypes. **B**, Number of synapses formed by each MG cell onto other classified cell types in the full EM volume (n = 5627 total synapses from 117 MG cells onto 310 categorized postsynaptic neurons; 7661 total synapses were annotated, with 27% unclassified). MG cells are ordered along the horizontal axis by the number of synapses onto ON (decreasing) and then OFF (increasing) cells. Across the MG population, there were 65 +/- 30 (sd) annotated synapses per cell. Pie charts illustrate the fraction (and total number) of synapses to each cell type from all MG cells represented in the bar graphs. **C**, Adjacency matrix showing the pairwise connection strength (number of synapses) for all MG and Output cells. **D**, Left: ON and OFF output cells for which presynaptic MG cells are shown. Middle: all presynaptic MG cells synapsing onto the Output ON cell (n = 18 MG- cells and 109 total synapses). Right: all presynaptic MG cells synapsing onto the Output cell OFF cell (n = 21 MG+ cells and 121 total synapses). **E**, Left: MG+ and MG- cells for which presynaptic MG cells are shown. Middle: all MG cells presynaptic to the MG+ cell (n = 13 MG- cells and 31 total synapses). Right: all MG cells presynaptic to the MG- cell (n = 13 MG- cells and 31 total synapses). **F**, Quartiles of the distribution for MG:Output convergence and divergence datasets (the whiskers extend to show the full distribution excluding outliers illustrated with open circles). **G**, Mean (and 95% ci) connection probability as a function of distance for MG cells synapses onto Output ON, OFF, MG+, and MG- cells. **H**, Total number of synapses from MG cells onto specific cellular compartments of Output (top) and MG (bottom) cells. **I**, Number of synapses from MG cells onto ON and OFF cells as a function of depth from the molecular layer. **J**, Number of synapses from MG cells onto MG+ and MG- cells as a function of depth from the molecular layer.

We also examined the convergence of MG cells onto Output and other MG cells and found a median of 8 MGs converging onto one Output cell and a 4:1 convergence ratio for MGs onto each other (Figures 5F and Extended Data Figure 4). As far as divergence, an MG cell synapses onto 6 Output cells and 5 other MGs (median values; Figures 5F and Extended Data Figure 4). These are likely underestimates due to additional MG cell inputs to (and outputs from) cells located outside the reconstructed volume. Connection probability declined with distance, consistent with synaptically connected MG and Output cells sharing electrosensory input from the same region of the body surface (Figure 5G).

### From synaptic connectivity to sensory prediction

To explore the functional roles of the different cell types and synaptic connectivity patterns revealed by our connectome analysis, we constructed a model that incorporates this synaptic connectivity and uses measured EAF and GCA responses and measured synaptic plasticity at GCA synapses onto MG and Output cells (Methods). The model is further constrained by matching recordings from MG and Output cells ^18^. Because little is known about the physiological properties of SG cells, we omit them from our model except for a study of their effect in Extended Data Figure 5.

As outline above, the ELL cancels the predictable self-generated component of the electrosensory input stream to reveal unpredictable signals of ethological relevance. This occurs in two ways: cell autonomous and multilayer. In cell autonomous cancellation, which occurs within Output cells, a "negative image" or inverse of the predictable input is extracted from GCA activity through synaptic plasticity. MG cells support cancellation through a second, multi-layer mechanism. MG cells construct a negative image from their granule cell input in a similar way as Output cells, but they transmit this as a prediction to the Output cells where it provides a second source of cancellation.

Learning of negative images in both Output and MG cells occurs through a plasticity mechanism that combines associative synaptic depression with a non-associative potentiation dependent solely on presynaptic spiking in GCAs ^14,35^. The overall effect of these two forms of plasticity is to drive the neuron toward a constant "equilibrium" rate of firing determined by parameters of the plasticity rule ^15,16^. If the sensory input to the neuron is total predictable, the result is that the plasticity-tuned GCA input precisely cancels the sensory input, leaving only a constant baseline input that drives firing at the prescribed equilibrium rate. When unpredictable inputs are present, the plasticity mechanism cannot achieve full cancellation and responses to the unpredictable input components remain. In the case of prey signals, this is exactly what the ELL is designed to achieve but, as we show in the next section, when noise is present partial cancellation can cause problems.

An important distinction between Output and MG cells is that plasticity in Output cells is driven by-and cancellation applies to-ordinary axonal action potential firing. For MG cells, plasticity is driven by dendritic spikes (called broad spikes), and cancellation fixes the rate of broad spike firing. Importantly, plasticity does not fix the rate of axonal (narrow) spike firing in MG cells, which allows them to transmit a prediction of the EOD responses to Output cells. As mentioned above, this separation of learning and signal transmission functions in MG cells requires special physiology, including the observed di-synaptic sensory input pathway.

To test our connectome / physiologically based model and to set some of its parameters, we simulated prior *in vivo* experiments (see Methods) in which cancellation can be monitored. In these experiments, the naturally occurring EOD is blocked by a paralytic, although EOD commands continue to be generated and an EOD mimic is triggered on these commands, providing direct experimental control over the relationship between EOD commands and their sensory consequences. Initially, there is no EOD mimic, and the ELL unlearns its prediction of the sensory consequences of the naturally occurring EOD. Then, when the EOD mimic is turned on, the learning process that leads to EOD sensory cancellation can be observed by examining the broad spike response to the EOD mimic in MG cells or the spiking response in Output cells. Under these conditions, both model and data show a rapid cancellation of the effects of the EOD mimic on broad spike firing in MG+ cells (n = 24) and a substantially slower cancellation of the spiking response of Output ON cells (n = 31; Figure 6A,B). MG- (n = 37) and OFF (n = 46) cells show little change in spike rate over 4 minutes of pairing (Extended Data Figure 5A-B) due to rectification (which conceals subthreshold changes), but intracellular recordings of Output OFF cells (n = 13) show ∼1 mV decay over 4 minutes of pairing from an initial response of ∼2.25 mV (Figure 6C). The time-course of cancellation in the model closely matches the data in all cases (Figure 6A-C; Extended Data Figure 5).

**Figure 6.**
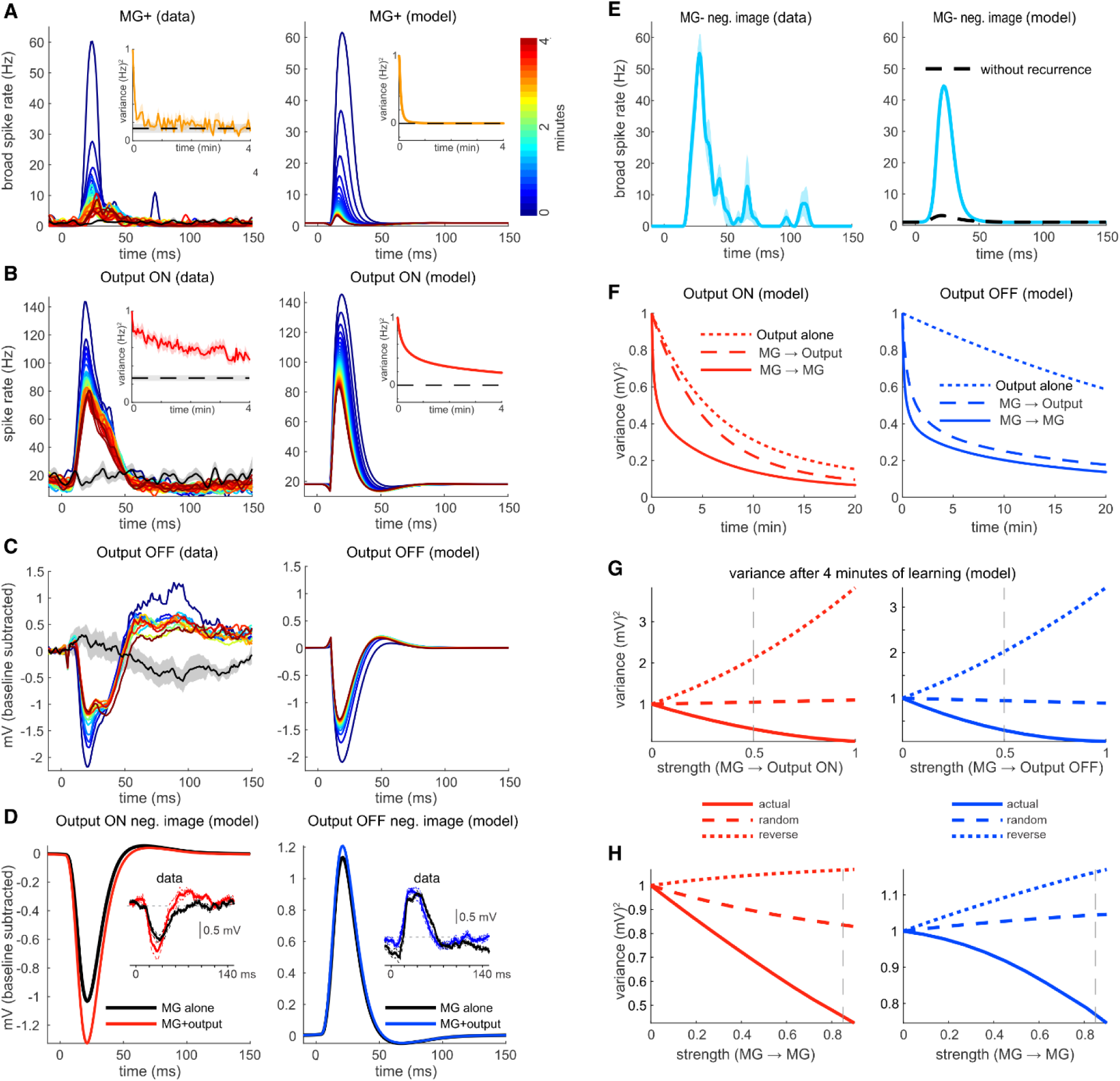
Effects of synaptic connectivity on sensory cancellation. A, Time-course of cancellation of the effects of an EOD mimic on the broad spike rate of MG+ cells for the data (left) and model (right). Time 0 on the x-axis in Figures 6-7 and Extended Data Figures 5&7 denotes the time of the EOD command. Lines are equally spaced from the start (blue) to the end (red) of the EOD-on period for both data and model. Black traces are the mean±SEM of the response before the EOD mimic is turned on. Insets show the time course of cancellation (mean±SEM) where lower variance indicates more accurate cancellation, and the dashed line shows the pre-pairing variance (mean±SEM). **B,** Time-course of cancellation for the spike rate in Output ON cells for the data (left) and model (right). **C,** Time-course of cancellation for the intracellularly recorded membrane potential of an Output OFF cells in the data (left) and model (right). **D**, Model Output cell membrane potential with the EOD-mimic turned off, revealing the learned negative image. For Output ON cells, ∼90% of the negative image is due to plasticity at GCA-MG synapses. The percentage is even higher for Output OFF cells, consistent with prior experimental findings^18^ (insets). **E**, Negative image in MG- cells in the data (left) and model (right). Black trace shows the negative image without MG-MG recurrence, indicating that very little of the learning is intrinsic to the MG- cell itself. **F,** Time-course of cancellation in the model as measured by the variance of the membrane potential, simulated for 3 conditions: (dotted) with learning only at GCA-Output synapses, (dashed) with adding learning at GCA-MG synapses but without MG recurrence and (solid) with MG recurrence. **G,H,** Variance of Output cell voltage responses after 4 minutes of exposure to the EOD mimic as a function of connectivity parameters for the same conditions as in **E**. Vertical dashed lines denote the default values in the model.

The different learning rates seen for Output and MG cells in Figure 6A,B is not the result of different learning rate parameters for these two cell types (they are the same). Rather, the difference arises in part from the larger number of GCA inputs that MG cells receive relative to Output cells (Figure 2A). In addition, the rate of depression of GCA synapses is proportional to the fractional change in firing rate induced by the signal being learned. Because the baseline rate for MG cells firing broad spikes is much lower than the Output cell baseline firing rate, the initial EOD sensory input increases the MG broad spike rate by a factor of ∼60, while the Output cell firing range only changes by a factor of ∼8. This is the primary reason that Output cell learning is slower than MG learning. In fact, in the result showed in Figure 6B,C most of the cancelation is coming not from Output cell learning, but from MG cell inhibition of the Output cell. To see this experimentally, the EOD mimic is turned off after the 4 min learning period, thereby removing the sensory input. This leaves only the learned GCA input which is the negative image, and in both the data^18^ and the model most (Output ON) or almost all (Output OFF) of the negative image comes from MG input (Figure 6D).

The effect of the change in broad spike rate on learning is different for MG+ and MG- cells. Whereas an MG+ cell can increase its broad spike rate by a factor of 60 due to the strong depolarizing effect of the EOD-related sensory input, the inverse of this signal can, at most, change the rate of broad spike firing in MG- cells from its baseline to 0, a factor of 1. This causes a dramatic disparity between the learning rates of MG+ and MG- cells. The negative image in MG- cells, seen by the same elimination of the EOD mimic as in Figure 6D, is well matched between the data and the model (Figure 5E), but in the model we can verify that almost all of this learned negative image is not arising from modification of GCA synapses onto the MG- cell, but rather it is transmitted from MG+ cells through recurrent inhibition (Figure 5E). Thus, in the learning of a positive going predictable sensory input, most of the learning is due to MG+ cells, with cancellation in MG- cells arising from their recurrent synaptic partners. For a negative going predictable sensory input, the situation would be reversed. An essential rationale for recurrent connections between MG cells is thus to remove what would otherwise by unavoidable learning asymmetries.

Having confirmed a close match between the model and data, we can further explore how cancellation performance depends on observed patterns of synaptic connectivity. MG cell inhibition of Output cells makes cancellation faster and more symmetrical between Output ON and OFF cells compared to a circuit with a single site of plasticity at GCA synapses onto Output cells (Figure 6F). Adding recurrent connections between MG cell subtypes further increases cancellation speed and symmetry in Output cells (Figure 6F, Extended Data Figure 5H). These benefits depend not only on the strength of MG connections, but critically, on the pattern of MG connectivity. Random connectivity between MG and Output cell subtypes confers no benefits in terms of cancellation speed, whereas reversed connectivity (i.e. a pattern opposite to that observed in the connectome) makes cancellation slower (Figure 6G). A similar pattern of results is observed for MG recurrent connections (Figure 6H).

### Multiple sites and timescales of plasticity support continual learning

The ELL uses at least two sites of synaptic plasticity for predictive learning and, as shown in the previous section, the learning rate at these two sites is quite different. What is the purpose of cancelling sensory input in two stages, instead of relying on a single site and what is the purpose of having two learning timescales? These questions are underscored by the connectome results demonstrating the circuit complexity and wiring specificity required to support a two-stage cancellation mechanism. To address this, we considered a more realistic scenario in which predictions must be maintained and updated in a dynamic environment in the presence of environmental noise. The fish we are studying are social and often search for prey surrounded by nearby conspecifics discharging their own electric organs ^36^. We simulated this situation using the model described above by switching between two contexts: a *self* condition (similar to the EOD-on condition in Figure 6) in which the circuit cancels a predictable EOD-like stimulus, and a *self + noise* condition in which the same EOD-like stimulus is combined with additional EOD-like signals with random amplitudes and polarities emitted at random times (Figure 7A; Methods). Cancellation performance was evaluated following a switch from the *self + noise* back to the *self* condition. Although these "noise" signals are unpredictable, they still affect the ELL learning system because synaptic plasticity at GCA synapses is always operational (i.e. this is a continual learning system).

**Figure 7.**
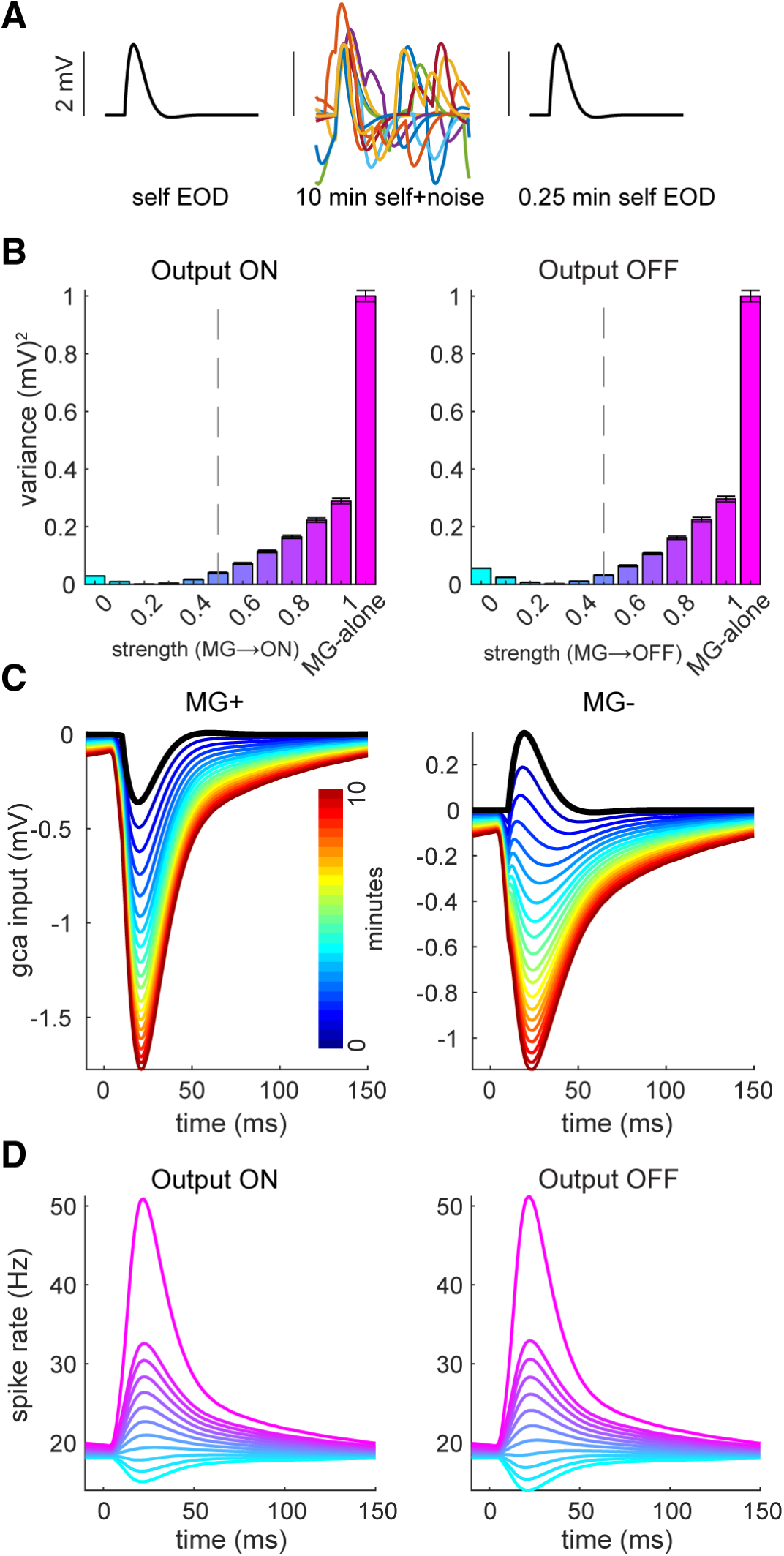
Multiple sites of plasticity solve a continual learning problem. A, Schematic of the simulation. Initially all variants of the model perfectly cancel a self-generated EOD signal. Then, random EOD-like noise is added for 10 min followed by a return to the self EOD context for 0.25 minutes. **B,** Average variance over the first 0.25 minutes following the switch to the self EOD context as a function of the strength of MG-to-Output cell inhibition. MG-alone corresponds to no learning in Output cells and with MG->Output coupling = 1. Averaged is over 100 simulations with different random noise. Error bars are SEM. **C,** Change in the GCA input to MG+ and MG- cells across 10 min of noise. Both cells start with the input needed to perfectly cancel the self EOD-like signal (black trace) but, in the presence of noise, both MG subtypes depress their GCA synapses. **D,** Spike rate of Output ON (left) and OFF (right) responses 0.25 min after the switch to the self EOD case. The line-colors in **D** correspond to the bar-colors in **B**.

Models that only have fast (MG-mediated) plasticity are severely affected by noise, so they perform poorly upon returning to the *self* context (Figure 7B; MG-alone). This occurs especially for the noise polarity that excites electroreceptors and arises from the asymmetry in learning rates between positive and negative sensory responses mentioned above. ‘Excitatory’ noise induces a rapid depression of GCA synapses that outweighs the slower synaptic potentiation induced by ‘inhibitory’ noise, leading to depression of GCA synapses onto MG cells (Figure 7C). This results in dis-inhibition of Output On and OFF cells (Figure 7D). In contrast, models with both fast (MG-mediated) and slow (Output cell-mediated) plasticity perform much better. Plasticity in Output cells mitigates the effects of the noise because the strong depression of GCA synapses onto MG cells dis-inhibits the output cells which in turn drives depression of their own GCA input, effectively cancelling the deleterious MG cell input. Combining the results shown in Figures 6 and 7, we see that the speed of learning of a predictable sensory input is enhanced by strong MG-to-Output inhibition, whereas robustness to noise favors weaker MG-to-Output inhibition. The value that we found best fits the data in Figure 6 appears to significantly speed learning while showing robustness to noise.

## Discussion

Our connectome analysis provides strong confirmation for a prior two-stage model of sensory prediction and cancellation in the ELL based on *in vivo* recordings ^18,19^. According to this model, the ELL faces two problems common to multi-layer learning systems ^37–39^. First, activity patterns required for generating predictions do not match those required for transmitting these predictions. In other words, the requirements for learning and signaling within individual neurons conflict. A potential solution to the first problem, based on synaptic inhibition and dis- inhibition, was identified previously from *in vivo* recordings and biophysical modeling ^18,19^.

Connectome analysis suggests that this solution is indeed implemented by the ELL. Second, the ELL faces a structural credit assignment problem—i.e. specific patterns of synaptic connectivity are required for correctly routing predictions from intermediate to output processing stages. A dense reconstruction of the synapses between MG and Output cell sub-types revealed remarkably precise connectivity matching that required for the cancellation of predictable sensory input.

Connectome analysis also revealed unexpected features of ELL circuitry, the functions of which remain to be explored. SG cells, for example, represent an additional prominent site of integration of electrosensory and predictive information. Because *in vivo* recordings from morphologically identified SG cells have not been made, we do not know whether their GCA inputs exhibit synaptic plasticity, as shown previously for MG and Output cells. If such plasticity does exist, our model shows that despite receiving fewer GCA inputs on an individual cell basis (Figure 2), SG cells can still contribute to the cancellation process (Extended Data Figure 6).

While the highly selective feedforward and recurrent connectivity of MG cells is well-suited to make cancellation of predictable input faster and more symmetrical between ON and OFF pathways, our model also suggests that MG cells must meet certain biophysical requirements in order to fulfill this role. Specifically, sensory-evoked inhibition (dis-inhibition) must selectively modulate dendritic spike firing, while recurrent inhibition must modulate both dendritic and axonal spike rates (Extended Data Figure 5I,J). While *in vivo* recordings have shown that the first requirement is met ^18,19^, the second remains to be tested. Intriguingly, differences in synaptic placement onto MG cells may explain how these biophysical requirements are fulfilled. Prior *in vivo* recordings and multi-compartmental modeling of MG cells showed that dendritic spike rate is substantially more sensitive to inhibition than axonal spike rate ^18,19^. SG synapses are mainly formed onto distal portions of MG basilar dendrites, electrotonically distant from the sites of axonal spike generation (Extended Data Figure 7). This location may be ideal for selectively modulating dendritic spikes. In contrast, recurrent MG synapses are located more proximally, closer to the sites of axonal and dendritic spike initiation, consistent with their impacting both spike types (Extended Data Figure 7). Additional electrophysiological studies, including *in vitro* recordings from synaptically coupled MG cells, in which effects of recurrent inhibition can be directly measured, are needed to test this and understand what additional roles SP- and Gr+ cells may perform.

The existence of multiple sites and forms of synaptic plasticity is a ubiquitous, yet poorly understood, feature of neural circuits, including regions associated with predictive processing such as the cerebellum, cerebral cortex, and hippocampus ^40–43^. In the case of the ELL, it has been unclear why the complicated, two-stage process posited previously and confirmed here is needed when a single site of fast plasticity at granule cell synapses onto Output cells would appear to be sufficient for cancelling predictable input. The connectome-constrained model developed here provides an answer: under realistic scenarios in which predictions must be maintained in the presence of noise, cancellation performance based on a single site of fast plasticity is poor due to ‘overfitting’ of the noise (Figure 7). While the idea that dual fast and slow learning systems could solve challenging continual learning problems is not new ^44–46^, the present work provides the actual implementation details of such a solution at the level of specific cell types and synaptic connectivity patterns.

There are several ways in which our connectome analysis of the ELL remains incomplete. While a high-degree of structure is clearly present in the connectivity between morphologically identified cell types, a substantial fraction of annotated synapses remained unclassified—either belonging to a cell within the volume that was not classified based on morphology or came from outside the volume. Additional inputs to the ELL not considered here include a direct feedback projection from a higher stage of electrosensory processing ^11,47^, ‘lateral’ connections from other parts of the ELL body map ^21^, and neuromodulatory inputs, including serotoninergic and noradrenergic fibers ^48,49^. Finally, the ELL molecular layer remains to be fully mapped. Key questions include the connectivity of molecular layer interneurons, the extent to which synaptically-coupled MG and Output cells receive shared versus distinct GCA input, and whether biases exist in patterns of GCA input onto ON and OFF MG and Output cell sub-types. The latter might be expected if learned predictions in the ELL leave a lasting anatomical mark.

How nervous systems generate predictive models of the interactions between an organism and its environment is a central question in neuroscience. The present study provides a detailed set of answers to this question by applying connectomics approaches to a uniquely tractable system in which sensory prediction has been studied for nearly 50 years. Given the many similarities between the circuitry, physiology, and plasticity of the ELL and other brain regions-- including the cerebellum and other cerebellum-like circuits ^33^, the hippocampus ^42,50^, and the cerebral cortex ^9,51^--we believe that these results provide a blueprint for understanding predictive processing in the vertebrate brain.

## Acknowledgments

This work was supported by grants from the National Institutes of Health (NIH NS075023 and NS118448) to N.B.S. and (U24NS109102, U19 NS104653, and UM1NS132250) to J.W.L. and from the Gatsby Foundation to LFA. Thanks to R. Andon, M. Chklovskii, D. Friedman, H. Hailu, R. Loike, S. Sasson, J. Singh, and A. Vaisse for assistance with neuronal reconstructions.

## Author contributions

S.Z.M, L.F.A., and N.B.S designed the research. M.P., R.S., Y.W., J.W.L., generated the EM volume. M.J. segmented EM volume. K.E.P., M.P., M.G. and N.B.S. analyzed the EM data. S.Z.M, A.G. and L.F.A. performed the computational modeling. S.Z.M analyzed the electrophysiological data. N.B.S, J.W.L, V.J, and L.F.A organized the project.

## Declaration of interests

The authors declare no competing interests.

## Data and code availability

EM image data and segmentation data will be available in Neuroglancer precomputed format will be made publicly available prior to publication. Python scripts for analysis of the data are available from the Github repository https://github.com/neurologic/efish_em. At the time of publication, supporting files (such as processed data), in .zip or original format, for individual scripts will be included in the Github repository, or if large, will be deposited on G-Node, as indicated in the script in question. Several scripts use Google Cloud BigQuery databases which are not publicly available and require a credentials file to access. However, the data contained within these databases will be freely available for download from the released data page.

## Methods

### Experimental model and subject details

Wild caught mormyrid fish (7-12 cm in length) of undetermined sex of the species *Gnathonemus petersii* were used in these experiments. Fish were housed in 60 gallon tanks in groups of 5-20. Water conductivity was maintained between 40-65 microsiemens both in the fish’s home tanks and during experiments. All experiments performed in this study adhere to the American Physiological Society’s *Guiding Principles in the Care and Use of Animals* and were approved by the Institutional Animal Care and Use Committee of Columbia University.

### EM data acquisition and processing

Two fish of the species *Gnathonemus petersii* were deeply anesthetized with tricaine methanesulfonate (MS:222, 1:10,000) and transcardially perfused with 2% paraformaldehyde and 2% glutaraldehyde in 0.15 M sodium cacodylate buffer. Following perfusion, 2.5 mm x 2 mm x 0.4 mm volumes of the hindbrain containing the anterior ELL were processed for electron microscopy using a modified rOTO staining protocol ^52^. The size of the skin region projecting to the imaged volume cannot be precisely estimated due to differences in receptor density and central magnification factor associated with different body locations. However, given its anterior location, the volume likely contains the central representation of a few mm of the skin surface on the chin appendage. The specimen ultimately selected for imaging was 9.3 cm long and weighed 6.9 g. After dissection, the tissue was washed three times for 10 minutes each in 0.15 M Sodium Cacodylate buffer (Sigma-Aldrich, C0250) at room temperature, then incubated in 2% osmium tetroxide (Electron Microscopy Sciences, #19190) in 0.15 M Sodium Cacodylate for 30 minutes flat followed by 4 hours on a rotator at room temperature. The sample was rinsed (3 × 10 min, 0.15 M Sodium Cacodylate) and incubated in 2.5% potassium ferrocyanide (Sigma-Aldrich, P3289) in 0.15 M Sodium Cacodylate for 2 hours at room temperature, followed by overnight incubation at 4 °C. The next day, samples were washed with Milli-Q water (3 × 10 min) and incubated in freshly prepared, 0.45 µm-filtered 1% (w/v) thiocarbohydrazide (Sigma-Aldrich, 223220) in water for 1 hour at room temperature. Following water washes (3 × 10 min), the tissue was post-fixed with 2% osmium tetroxide (Electron Microscopy Sciences, #19190) in water for 4 hours at room temperature, washed again (3 × 10 min), and incubated overnight at 4 °C in 2% aqueous uranyl acetate (Electron Microscopy Sciences, #22400). Samples were brought to room temperature the following day, placed in a 60 °C oven for 1 hour, and returned to room temperature before final water washes (3 × 10 min). Dehydration was carried out in a graded ethanol series (30%, 50%, 75%, 95%, 3 × 100%) using wide-mouth pipettes to avoid tissue damage. The tissue was infiltrated with increasing concentrations of LX-112 resin in propylene oxide (25%, 50%, 75%) and then with 100% resin, freshly prepared using 29.1 g NMA, 46.8 g LX-112, 14.1 g NSA, and 1.8 g BDMA (Ladd Research, #21212) Resin infiltration was performed over 2 days with uncapped tubes to allow evaporation of propylene oxide. On the final day, the sample was embedded in BEEM capsule molds and cured at 60 °C.

The resin embedded sample was trimmed and sectioned using an ultramicrotome (UC6, Leica, Germany) fitted with Diatome Trim 90 and Diatome Ultra 45/Ultra 35 diamond knives, respectively (Diatome, USA). A 250 µm depth of tissue was serially sectioned at ∼30 nm thickness using an automated tape-collecting ultramicrotome (ATUM) system ^53^ and collected onto carbon-coated Kapton tape. Sections were mounted 120 per wafer across 69 silicon wafers using double-sided carbon tape. Wafers were post-stained with uranyl acetate and lead citrate prior to imaging with a multibeam scanning electron microscope (MultiSEM, Zeiss) as in ^29^. Wafer overviews were first acquired using reflected light microscopy (Zeiss Axio Imager) to guide automated acquisition. Imaging proceeded in phases using a 61- beam configuration with 4 nm pixel size and 200ns dwell time. A custom workflow manager monitored acquisition, validated data integrity and focus, and generated real-time diagnostic reports (https://github.com/YuelongWu/mSEM_workflow_manager). We collected images spanning a 105 µm depth, totaling 3,534 sections, of which 83 non-consecutive sections were missing with the exception of one instance with three consecutively skipped sections. Stitching was performed using a block-matching algorithm on a scattered grid within the overlapping regions of image tiles, followed by elastic optimization across high-resolution tiles. The stitched images were then rendered into non-overlapping 4096×4096 px tiles. To enable large-scale registration, low-resolution thumbnail images (512nm pixel resolution) were generated in parallel with acquisition and aligned using a combination of feature-based affine registration, template matching, and spring-mesh relaxation. The transformations resulting from this thumbnail alignment were then upscaled and used as an initial estimation for fine alignment. The fine alignment process involved its own template matching and spring-mesh relaxation steps, but at a much higher resolution (16nm) compared to the thumbnails. Finally, the resulting alignment transformations were applied to the full-resolution stitched dataset. The stitching and the alignment processes were carried out with a combination of feabas (https://github.com/YuelongWu/feabas) and mb_aligner (https://github.com/adisuissa/mb_aligner).

### EM segmentation and skeletonization

We first applied 16 nm and 32 nm Flood-Filling Networks (FFNs), originally trained on a human cortical dataset ^29,30^, to the ELL dataset to generate a preliminary segmentation. This preliminary output was manually agglomerated to create a ground truth dataset consisting of complete axons and dendrites. Using this ground truth, we retrained the original human cortex FFNs, and used sparse point pair annotations to select optimal model checkpoints. This yielded optimized 16 nm and 32 nm FFN models, which were used to produce a base segmentation tuned for low merge error and an agglomeration that reduced split errors. At the agglomeration stage, we applied soma splitting using 12,700 manually annotated soma using VAST ^54^. Skeletons were then generated from the base segmentation using the same TEASAR-based method employed for the human cortical dataset.

### Cellular Reconstruction

After base segmentation and agglomeration of the EM volume was completed, a modified version of CREST was used to reconstruct all neurons and neuron fragments in the current dataset ^29^. Complete instructions and a demonstration of the proofreading process in CREST is available via the CREST homepage (https://github.com/ashapsoncoe/CREST). The modified CREST codebase (eCREST) used in this study is available via the eCREST homepage (https://github.com/neurologic/eCREST). When proofreading a biological object using CREST, a reconstruction was initiated from a seed base segment and all of the base segments that were agglomerated with it. Split errors were corrected by adding on missing base segments (and their agglomerated base segments) and merge errors were corrected by removing base segments that did not belong to the object (which was made more efficient by CREST). An object was considered complete once all of the base segments that define its morphology were identified, and no base segments that were not part of it were included. Correction of merge errors within an individual base segment was not possible. CREST enabled all base segments in each branch of a neuron to be labeled according to their subcellular compartment (axon, apical dendrite, basal dendrite, and soma). When a single base segment significantly spanned multiple structures, it was categorized as ‘multiple.’ For each proofread neuron object, the classified final set of base segments were recorded into JSON state which could be reloaded in eCREST for annotation (of synapses and soma), inspection and/or further proofreading.

### Morphological Identification

Cells that receive plastic granule cell input from the molecular layer are distinguished by spiny apical dendrites – these include two classes that have been characterized anatomically and electrophysiologically: Output cells and MG cells ^20–27^. These two classes have the largest soma of cells ELL, with output cells tending to be larger than MG cells. MG cells are clearly distinct from Output cells in that the former are interneurons, synapsing locally within the ELL, whereas the axons of Output cells exit the ELL without forming collaterals. The morphology of the basal dendrites of MG and Output cells are also highly distinct. A third class of cells with spiny apical dendrites has been previously identified morphologically are the SG cells ^21,25^. Like the larger MG cells, SG cells have spiny apical dendrites and axons that terminate locally within the ELL. Although it is possible that SG and MG cells form a continuum, for consistency with prior work and to aid the present analysis we objectively split these classes using hierarchical clustering (Ward method with Euclidean distance) based on the following morphological criteria: soma size, lateral spread of basal dendrite and axon (along x and z axes), and apical dendritic size (Extended Data Figure 2). Point annotation classes in neuroglancer (implemented via eCREST) were used to mark and measure the diameter and location of each neuron’s soma. We used the statistics of the x,y,z locations of the set of skeleton nodes (from the base segment skeletonization) for all base segments in each structure for each cell to quantify the basal dendrite and axon spread (std in x and y) and apical dendrite size (node count). There were 46/728 MG and SG cells that were manually identified as having either dendrites or axons out of the volume, or soma partially out of the volume could not be included in the hierarchical clustering; these cells were classified manually based on the remaining parts of the cells that were within the volume.

Output ON and OFF cell subtypes are distinguishable by soma positions in the plexiform and ganglion layers, respectively. Output ON cells also tend to have a more fusiform soma shape. We used the basal dendrite location of the MG+/MG- subtypes to assist in defining a boundary plane between the ganglion and plexiform layers (Extended Data Figure 2). Two Output cells had soma intersecting the ganglion- plexiform boundary and were categorized by their soma shape (fusiform versus round). The basal dendrites of MG+ cells arborize in the plexiform layer, while those of MG- cells arborize in the ganglion layer. The axons of MG+ and MG- cells have the opposite pattern: MG+ type arborize in the ganglion layer, while MG- type arborize predominantly in the plexiform layer (though they also send terminals into the ganglion layer). Therefore, the relationship between axon and basal dendrite position in these two subtypes is anti-correlated and could be used, along with a linear regression classifier, to completely split the MG population into MG+ and MG- types (Extended Data Figure 2). We extended the subtype morphological characterization of MG cells to SG cells as well according to the Type 1&2 morphological characterization previously established for MG cells (MG-:MG1 and MG+:MG2). The basal dendrites of SG- cells arborize in the ganglion layer (like MG-), while the basal dendrites of SG+ cells arborize in the plexiform layer (like MG+). Like MG cells, the axons of SG- and SG+ cells have the opposite layer pattern as their dendrites: SG- branch predominantly in the plexiform layer, while SG+ branch in the ganglion layer. Therefore, the relationship between axon and basal dendrite position in these two subtypes is also anti-correlated and could be used in a linear regression model to split the SG population into SG+ and SG- subtypes (Extended Data Figure 2).

Other cell types that we classified in this study are: the electroreceptor afferent fibers (EAF), the granular (Gr+) cell, and the small plexiform (SP-) cell (Extended Data Figure 2). Granular cells (Gr+) are the most numerous cell type in the ELL (aside from the granule cells of the overlying eminentia granularis posterior) ^20,21^. Despite their similar names, only the latter give rise to the granule cell axons in the molecular layer. Gr+ cells have small soma and their axon extends toward the plexiform and ganglion layers, branching throughout both. Basal dendrites of Gr+ ramify through the granular layer of ELL where EAFs terminate. If any apical dendrite exists, it is smooth and constrained to the granular layer as well. SP- cells have similarly small soma to Gr+ cells, but their soma are mostly in the plexiform layer. The axon of SP- cells, after exiting the soma, splits into two branches that go in opposite directions: one branch goes toward (and often into) the molecular layer with synaptic boutons terminating in both the ganglion and molecular layers, and another branch goes toward the granular layer with synaptic boutons terminating locally within the plexiform layer and sometimes in the granular layer. The apical dendrite of SP- cells is variable length (sometimes extending into the molecular layer and sometimes constrained to the plexiform layer), but is always smooth. Reconstructions that did not fit into one of the above categories, including neuronal processes for which no soma in the volume could be identified, were termed ‘unclassified.’

### Synapse identification and annotation

Following training in synapse identification from M.P., K.P., M.G., and N.B.S. manually identified and recorded synapses using neuroglancer/eCREST. Synapses were identifiable based on several key features: (i) the presence of a synaptic cleft, characterized by a thickened membrane gap at the junction between an axon bouton and a postsynaptic neuronal compartment, which is consistently observed across multiple tissue sections; and (ii) the presence of an *active zone*, indicated by the clustering of synaptic vesicles at the presynaptic membrane adjacent to the apposition site. Point annotation classes in neuroglancer (implemented via eCREST) were used to record the location of synapses to/from a reconstructed neuron. Presynaptic synapse annotations were placed on the presynaptic side of each identified synapse and on a base segment such that the base segment associated with that synapse would be recorded in the reconstruction file. Postsynaptic synapse annotations were placed on the postsynaptic side of each identified synapse. For each annotated synapse, the annotated base segment was either matched with an existing reconstruction containing that base segment, or a new reconstruction was seeded and completed from that base segment if it could not be found in any existing reconstruction. After all synapse base segments were accounted for in the dataset, we could create connectivity matrices among reconstructed cells.

### Conditional output/input analysis

We implemented the conditional input analysis methods described by ^55^ (see also ^56^). We constructed a connectivity matrix in which we counted the total synaptic connections between each cell and each pre/postsynaptic cell type (for conditional input/output analysis respectively). Conditional Input (associated with Figures 3, 4): On the condition that a cell gets one or more synapses from one cell type A, we counted the number of input synapses that cell gets from another cell type B (less the synapse from cell type A that qualifies the cell for each condition) for all possible pairs of cell types. Because not all synapse sets were fully annotated for each cell, we normalized the connectivity matrix for each cell to the fraction of total input from each cell type (number of synapses from each cell type divided by the total number of synapses across all cell types). Conditional Output (related to Extended Data Figure 4): On the condition that a cell makes one or more synapses to one cell type A, we counted the number of output synapses that cell makes to another cell type B (less the synapse from cell type A that qualifies the cell for each condition) for all possible pairs of cell types. Because not all axons were fully reconstructed for each MG cell (due to axons exiting the volume), we normalized the connectivity matrix for each cell to the fraction of total synapses to each cell type (number of synapses to each cell type divided by the total number of synapses across all cell types). Inputs/outputs across all cells are then compared to a randomized null model. In the null model, for each row-wise cell, each synapse is assigned to a randomly selected cell from among the existing column-wise set of cells (ie. cells that are already contained in the original connectivity matrix for each conditional analysis; which is a subset of the cells reconstructed for the whole study). In this way, the number of synapses each row-wise cell gets/makes remains constant, but the post-/pre-synaptic partner identities are randomized. We generated 100 random models of shuffled connectivity to generate a null distribution of conditional input/output counts. For each pair of inputs/outputs, we computed a z-score from this null distribution for the observed connectivity (i.e., how many standard deviations from the mean of the null distribution the observed number of counts is). The resultant conditional input/output matrix could be used to determine if observed connectivity from/to a specific cell type was significantly more or less dependent on connectivity from a second input/output than would be expected in a random model of connectivity. In the resultant conditional input/output matrix, the color scale for each cell in the matrix indicates the z-score. Cell types (conditions and inputs/outputs) were grouped along the axes according to their known/putative physiological responses to electrosensory stimuli.

### Electrophysiological recordings

Neural data presented in Figure 6 is a re-analysis of data collected in a prior study ^18^. Briefly, fish were anesthetized (MS:222, 1:25,000) and held against a foam pad. Skin on the dorsal surface of the head was removed and a long-lasting local anesthetic (0.75% Bupivacaine) was applied to the wound margins. A plastic rod was cemented to the anterior portion of the skull to secure the head. The posterior portion of the skull overlying the ELL was removed. Gallamine triethiodide (Flaxedil) was given at the end of the surgery (∼20 μg/cm of body length) and the anesthetic was removed. Aerated water was passed over the fish’s gills for respiration. Paralysis blocks the effect of electromotoneurons on the electric organ, preventing the EOD, but the motor command signal that would normally elicit an EOD continues to be emitted at a rate of 3 to 5 Hz. The EOD motor command signal (the synchronized volley of electromotoneurons that would normally elicit an EOD) was recorded with an electrode placed over the electric organ. Extracellular single-unit recordings were made from the VLZ using glass microelectrodes (2-10 MΩ) filled with 2M NaCl. For *in vivo* whole-cell recordings electrodes (8-15 MΩ) were filled with an internal solution containing, in mM: K-gluconate (122); KCl (7); HEPES (10); Na2GTP (0.4); MgATP (4); EGTA (0.5), and 0.5-1% biocytin (pH 7.2, 280-290 mOsm). No correction was made for liquid junction potentials. Only cells with stable membrane potentials more hyperpolarized than -45 mV and spike amplitudes >40 mV were analyzed. Membrane potentials were recorded and filtered at 3-10 kHz (Axoclamp 2B amplifier, Axon Instruments) and digitized at 20 kHz (CED micro1401 hardware and Spike2 software; Cambridge Electronics Design, Cambridge, UK). For pairing experiments shown in Figure 6, neural responses to the EOD motor command were recorded before, during, and after 4 minutes of *pairing*, in which an EOD mimic was presented 4.5 ms following EOD command onset. The EOD mimic was a 0.2 ms duration square pulse delivered between an electrode in the stomach and another positioned near the electric organ in the tail. The amplitude was 25 µA at the output of the stimulus isolation unit (stomach electrode negative).

### ELL model

As in prior models of the ELL, we modeled the response of the cells and learning discretely for each EOD (which we denote here with an index *i*). In the experimental setup we simulate the average EOD rate is 4 Hz, so a 4 min period of EOD learning corresponds to 4 min*60 s/min *4 EOD/s = 960 EODs.

We define *S* as the EOD-generated stimulus to be cancelled, which is a decaying sinusoidal function that we fit to the shape of the average EOD-evoked stimulus in the data. The amplitude of *S* is adjusted to produce a 2.25 mV change in the membrane potential to match the data (Figure 6). We track S and the different model activities for a time period from 10 ms before the EOD command to 150 ms after it in time steps of 1 ms. Thus, *S* is a *T* = 160 component vector across this time span. GCA activity in the model was taken from *in vivo* intracellular recordings of granule cell response to EOD corollary discharge input ^15^, and distributions of GCA synapses onto MG, Output and SG cells were obtained from the connectome. The GCA activity matrix is of dimension *T* x *N*, where *N* = 2840 for Output cells, *N* = 6390 for MG cells and, when SG cells are included, *N* = 246 for them. Except for Extended Data Figure 7 we did not include learning at GCA-SG synapses as the model shows minimal contribution to speed of cancellation from these synapses (see Discussion).

The model includes dis-inhibitory and inhibitory sensory input pathways, MG+, MG-, Output ON, and Output OFF cells. The baseline-subtracted membrane potentials of ON and OFF Output cells are *T*- component vectors determined at the time of the *i*^th^ EOD by

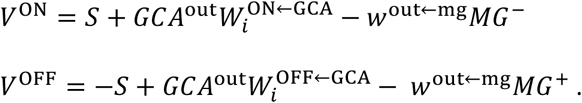

Here 𝐺𝐶𝐴^out^is the GCA activity (quantified as the probability of spiking at each time bin) synapsing onto Output cells convolved with a double exponential to yield an EPSP input. W^ON←GCA^ and W^OFF←GCA^ are the synaptic weights from the GCAs to the ON and OFF Output cell. They depend on *i* because of synaptic plasticity. 𝑀𝐺^±^ are *T*-component MG cell activities and 𝑤^out←mg^ describes the strength of inhibition from MG to Output cells. MG activities are delayed by 1 ms to account for the di-synaptic input-MG-Output pathway. For the reverse connectivity simulation in Figure 6H we exchanged the 𝑀𝐺^±^ subtype in both of the above equations, and for the random connectivity we added both MG inputs to both Output cell types weighted by 0.5𝑤^out←mg^.

𝑀𝐺^±^ represent the narrow spiking activity of MG cells, and are determined by a set of coupled equations

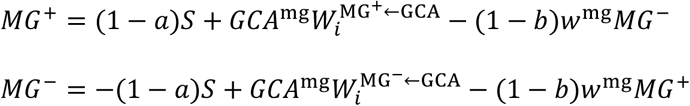

where 𝑎 and *b* parameterize the selectivity of narrow spikes to sensory and MG input, respectively, which is used for Extended data Figure 5I,J, but in all other cases, 𝑎 = 1 and 𝑏 = 0.

To compute the broad spiking activity of MG± cells, we first compute a membrane potential that controls broad spiking,

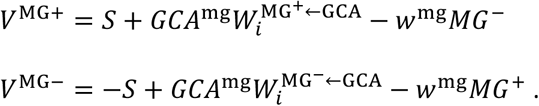

For the reverse connectivity simulation in Figure 6I, we exchanged the MG± subtypes in these equations, and for the random connectivity we added both MG inputs to both MG cell types weighted by 0.5𝑤^mg^.

For MG narrow spikes, which have a high baseline rate and are not typically subject to rectification, there is a linear relationship between membrane potential and firing rate^19^. However, for Output and MG broad spike activity, we transformed their membrane potentials into firing rates using a sigmoid nonlinearity^19^.

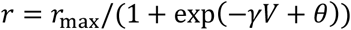

with 𝛾𝛾 = 1.59/mV, 𝜃𝜃= 2.20 and 𝑟_max_ = 200 Hz for Output cells, and 𝛾𝛾 = 2.53/mV, 𝜃𝜃= 4.30 and 𝑟_max_ = 75 Hz for MG cells (broad spikes). This sets the equilibrium rates (for *V* = 0) to 20 Hz for Output cells and 1 Hz for MG broad spikes, and the stimulus response to a 2.25 mV depolarization to 160 Hz for Output cells and 60 Hz for MG broad spikes.

Synaptic plasticity at GCA synapses onto MG and Output cells was based on prior *in vitro* and *in vivo* measurements ^13,14,36,37^. Plasticity in the ELL is anti-Hebbian with

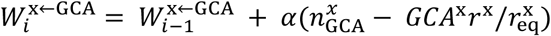

where x can be either "out" (Output cell) or "mg" (MG cell), and 𝑛^𝑥^ is a vector with components equal to the number of spikes fired by each GCA over the course of the *i*^th^ EOD. The parameter 𝛼 = 10^−9^mV sets the overall learning rate.

### Learning in the noise context (**Figure 7**)

In the *self + noise* condition, the noise consisted of ∼16 Hz EOD-like pulses delivered at random times (i.e. temporally uncorrelated with granule cell corollary discharge inputs). Noise amplitudes were drawn from a uniform distribution ranging from -1 to 1, where 1 corresponds to the self EOD and -1 corresponds to the same amplitude as the self EOD but opposite polarity. Opposite polarity EOD-like stimuli induce opposite polarity spike rate modulations in electroreceptor afferent nerve fibers. For the initial parameters we set the plastic weights so that the MG and Output cells canceled the self EOD signal perfectly.

### SG model (Extended Data Figure 6)

The separation of learning and signaling into dendritic and axonal spikes in MG cells enables MG cells to transmit a negative image. On the other hand, we assume that, even if SG cells generate two spike types, they are not sufficiently electronically separated to separate learning and signaling. Thus, unlike MG cells, SGs receive and transmit the sensory input and, if learning takes place at GCA-SG synapses, it will cancel the EOD-evoked sensory input at the level of axonal spikes. Thus, we assume that SG cells transmit a “cleaned up” version of the sensory input in which predictable features have been partially cancelled.

## Extended Data

**Extended Data Fig. 1.**
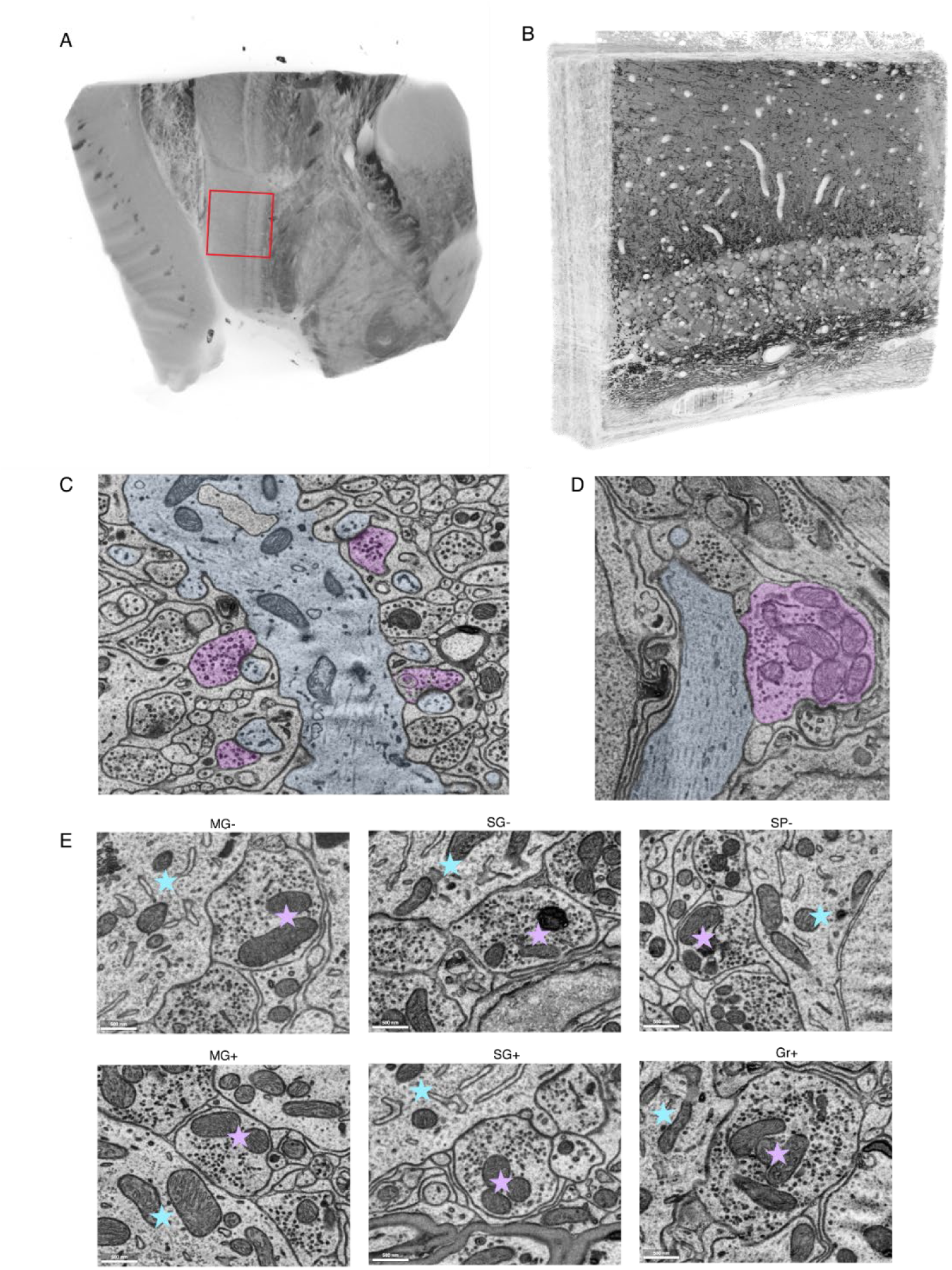
ELL sample preparation and EM data examples (Related to. Figure 1**). A,** uCT volume of a transverse section through the hindbrain containing the region of interest within the ventrolateral zone of the ELL (dashed rectangle). **B,** EM volume of the ELL obtained at 4 nm x 4nm x 30 nm resolution. **C,** Examples of asymmetric, putative excitatory, synapses onto apical dendritic spines in the molecular layer. **D,** Example symmetric, putative inhibitory synapse formed by an MG cell. In **C-D** dendrites are highlighted in light blue, axons in purple. **E**, Additional example of asymmetric synapses (purple stars) formed onto somata (blue stars) by different ELL cell types.

**Extended Data Fig. 2.**
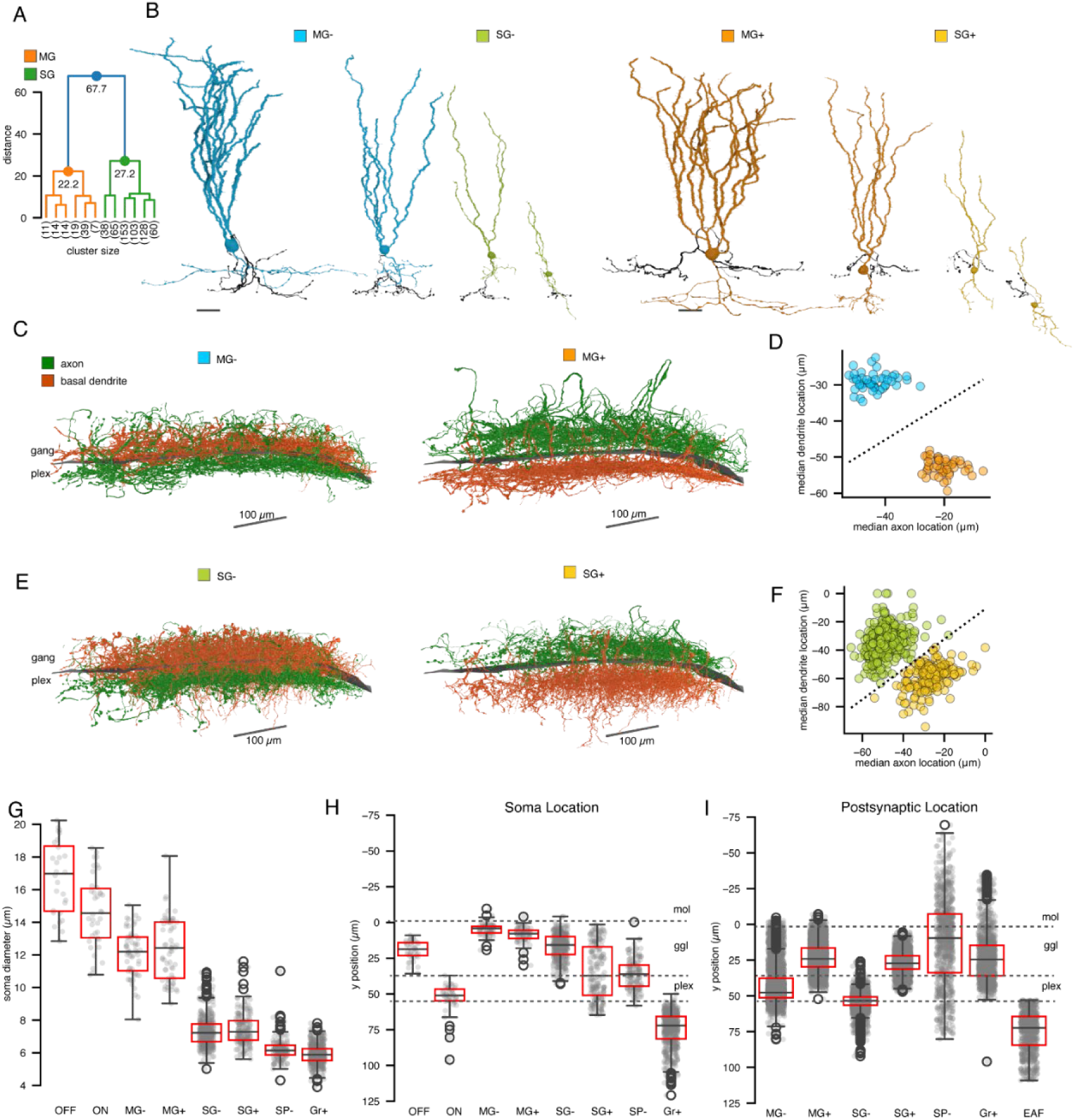
Morphological characterization of ELL cell types (Related to. Figure 1). A, Dendrogram visualization of the hierarchical clustering used to classify MG (n = 104) and SG (n = 547) cells. An additional 32 MG and 45 SG were manually categorized because they lacked sufficient soma in the volume and/or the majority of either their axon or dendrite exited the volume. B, Example MG and SG cells showing the variation in morphology across the population. Axons visualized in black. C, Reconstructed axons and basal dendrites of all MG cells showing the inverse relationship between axon and basal dendrite position (perpendicular to the layer plane in ELL) for MG- versus MG+ subtypes. **D**, Scatterplot of the median basal dendrite versus axon position for all MG cells. Dotted line shows the classifier obtained by linear regression on the dataset. Any cells shown in **C** with either axon or basal dendrite mostly out of the volume were excluded from this analysis and classified manually based on the location of the remaining in-volume axon/dendrite. **E**, Reconstructed axons and basal dendrites of all SG cells showing the inverse relationship between axon and dendrite position (perpendicular to the layer plane in ELL) for SG- versus SG+ subtypes. **F**, Scatterplot of the median basal dendrite versus axon position for all SG cells. Dotted line shows the classifier obtained by linear regression on the dataset. Based on these classification metrics, the SG- and SG+ subtypes are less distinct than MG+/MG- because of how much further ventrally SG- basal dendrites extend compared to MG-. Any cells shown in **E** with either axon or basal dendrite mostly out of the volume were excluded from this analysis and classified manually based on the location of the remaining in-volume axon/dendrite. **G**, Scatterplots (data points are randomly jittered along the x-axis within each category) overlaid on boxplots illustrating the quartiles of the distribution for soma diameter of each cell type (the whiskers extend to show the rest of the distribution excluding outliers illustrated with open circles). Only cells with their full soma in the volume were included in this analysis. **H**, Same display as **G** for the distribution for soma location of each cell type. **I**, Same display as **G** for the distribution of the locations of synapses formed by each cell type.

**Extended Data Fig. 3.**
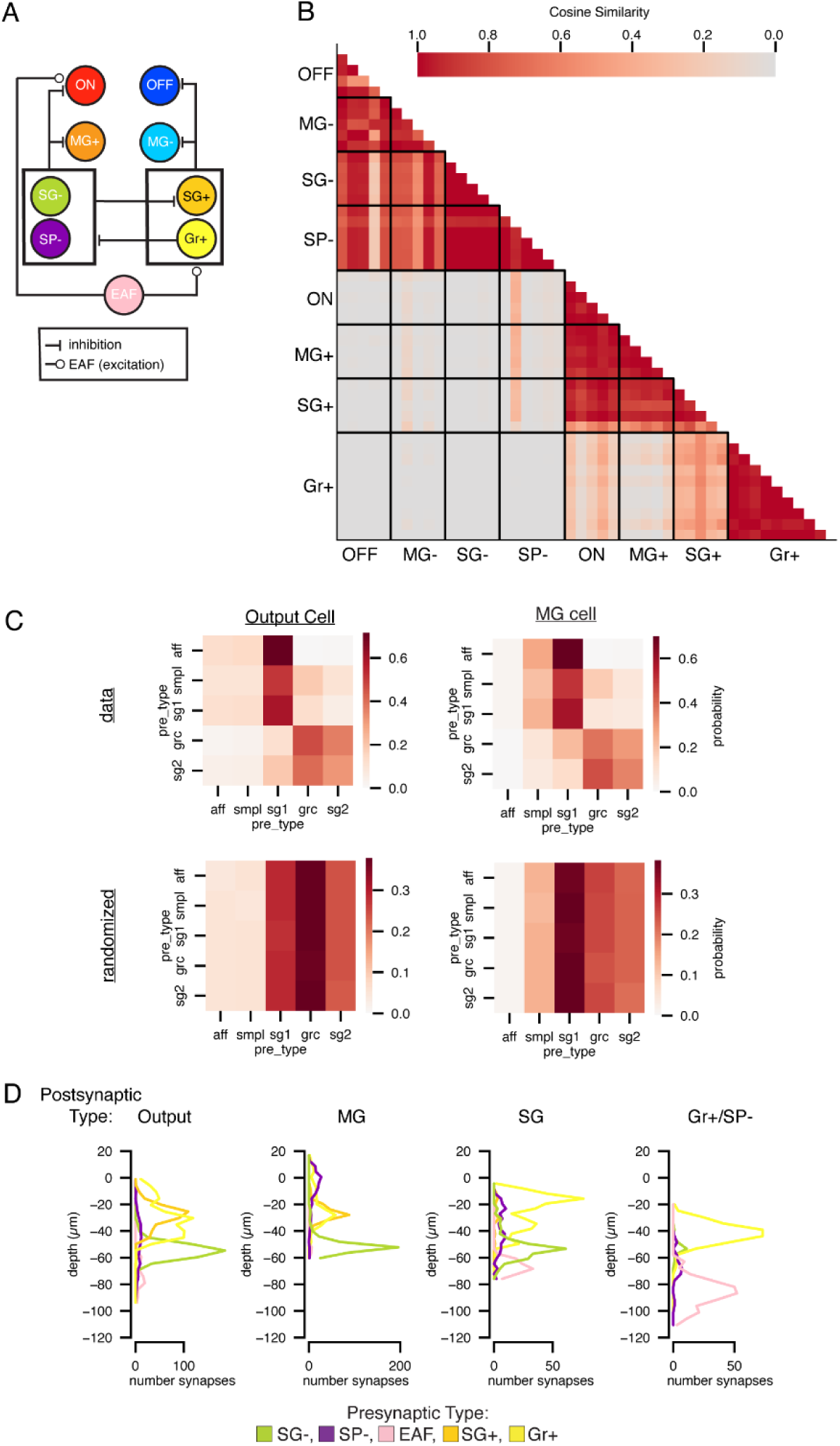
Sensory input pathways (related to. Figures 3 **& 4). A**, Summary diagram of the sensory input pathways from EAFs to Output and MG cells (only connections >10% of total synapses to each cell type are shown). **B**, Pairwise cosine similarity of the input pattern per cell type for all cells whose presynaptic sites were annotated and their inputs reconstructed. Cosine similarity was calculated on the total number of input synapses to each postsynaptic cell from each presynaptic cell type (after within-cell normalization to the total number of input synapses for each cell). Cells are ordered along the x and y axes according to their cell types, and cell types are ordered according to putative ON- versus OFF-type physiology (based on connectivity and/or *in-vivo* data from previous work). **C**, Data: The conditional input across Output and MG cells without z-scoring applied. The color of each square represents the mean fraction of input synapses from each cell type conditional on getting a synapse from a given cell type (condition). Randomized: The results of one iteration of the randomized data shuffle used for the null model of connectivity (see Methods). The color of each square represents the mean fraction of input synapses from each cell type conditional on getting a synapse from a given cell type (condition). Cell types are grouped along the x and y axes according to their putative sensory responses (inferred from connectivity and/or known from previous *in vivo* recordings). **D**, Histograms of presynaptic input location (relative to the ganglion-molecular layer boundary) for all inputs from each cell type. All cells used for the analyses in Figures 3 and 4 are included here.

**Extended Data Fig. 4.**
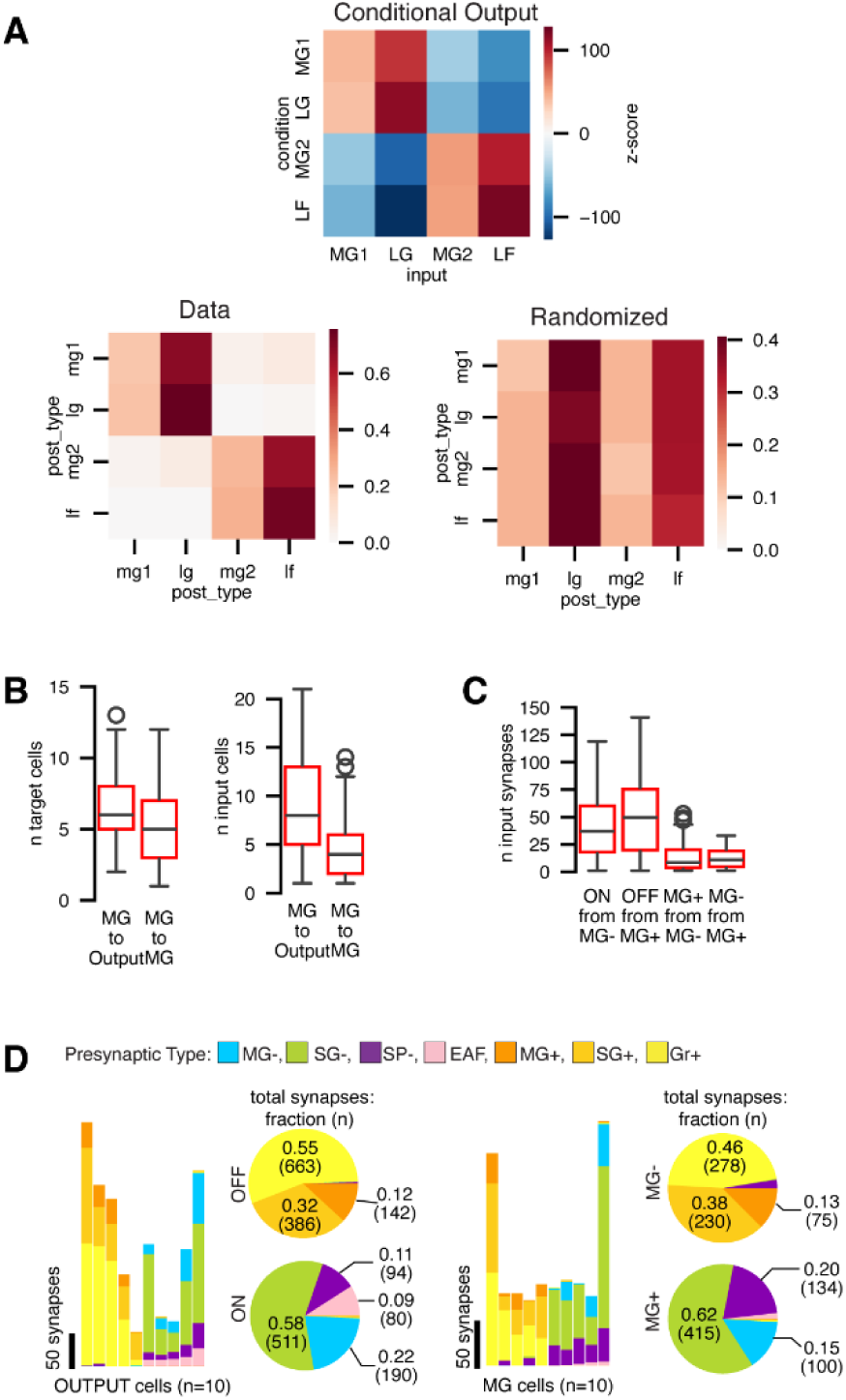
Feedforward and recurrent connectivity of MG cells (related to. Figure 5**). A**, Top: Observed MG cell connectivity with Output and other MG cells compared with random connectivity. Conditional input analysis was applied to the 117 MG cells represented in Figure 5. Postsynaptic cell types are grouped along the x and y axes according to their putative sensory responses (inferred from connectivity and/or known from previous *in vivo* recordings). Two clusters (illustrated with dotted lines) are observed among the input cell types, corresponding to the ON and OFF pathways. Bottom, Data: The conditional output across all MG cells without z-scoring applied. The color of each square represents the mean fraction of synapses onto each cell type conditional on synapsing onto a given cell type (condition). Randomized: The results of one iteration of the randomized data shuffle used for the null model of connectivity (see Methods). The color of each square represents the mean fraction of synapses onto each cell type conditional on synapsing onto a given cell type (condition). **B**, Divergence and convergence of MG connectivity to all Output cells (n = 117 presynaptic MG cells and 87 Output cells) and to/from all MG cells (n = 104 presynaptic and 116 postsynaptic MG cells).

**Extended data Figure 5.**
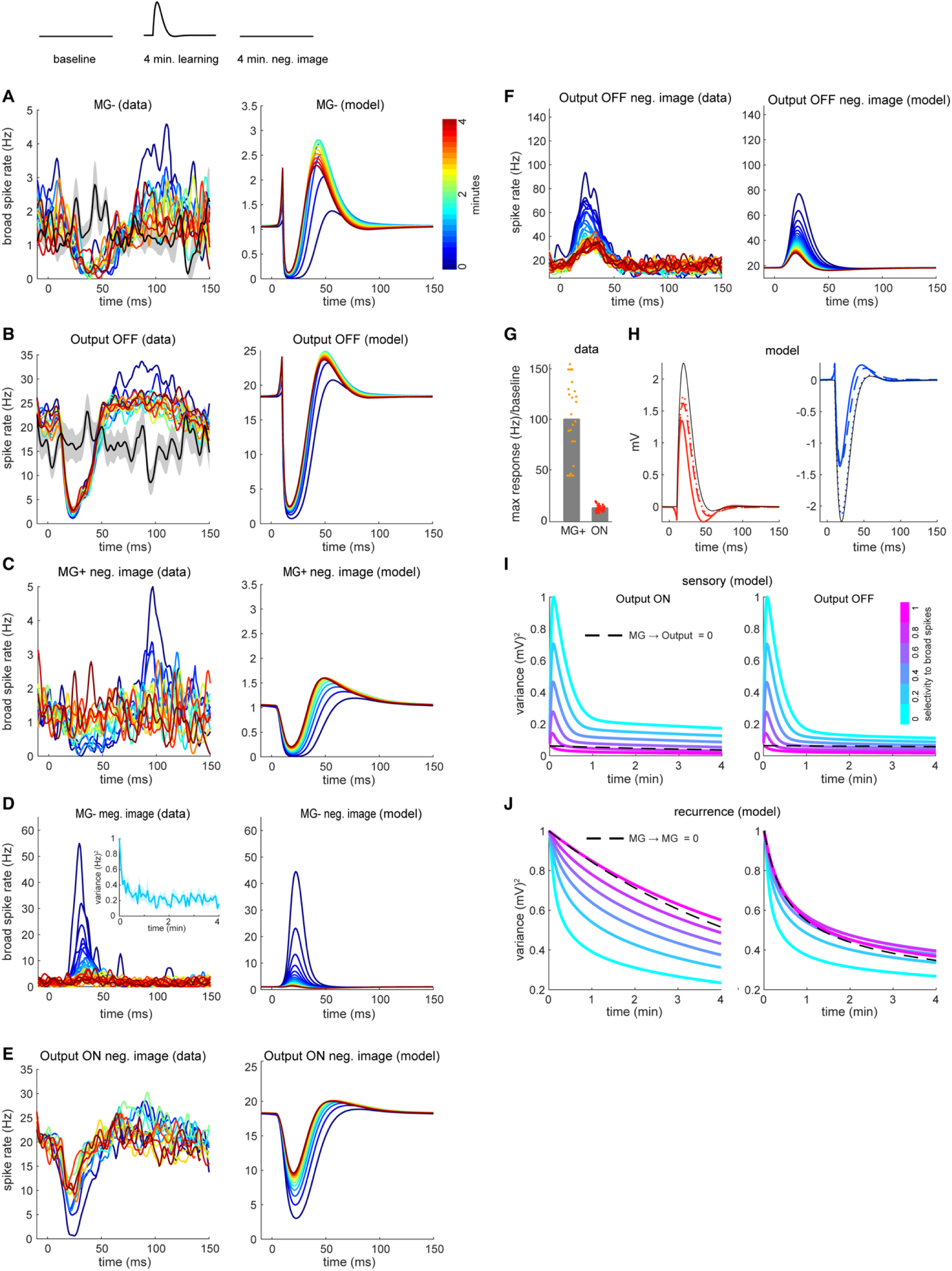
Effects of synaptic connectivity on sensory cancellation (related to. Figure 6). Top, schematic of the experimental design for measuring sensory cancellation. **A-B,** Time-course of cancellation of the effects of the EOD mimic on spike rate in MG- cells (**A**) and Output OFF (**B**) in the data (left) and model (right). Lines are equally spaced from the start (blue) to the end (red) of the EOD period. Black traces are the mean±SEM of the baseline response. **C-F,** Time-course of recovery of the negative image from effects of learning to cancel the EOD mimic on dendritic (broad) spike rate in MG+ cells (**C**), MG- cells (**D**), Output ON (**E**) and Output OFF (**F**) in the data (left) and model (right). **G,** Ratio of maximal rate response in the first second relative to baseline firing. On average, MG+ dendritic firing has a ratio of ∼100 and Output ON of ∼12 (in the model we matched the average response across all cells). **H**, Spike rate of Output ON (left) and Output OFF (right) after 4 minutes of learning in the model for the different conditions described in Figure 6F. The dotted line is Output alone, the dashed line is with MG-Output coupling but without MG recurrence and the full-line is with MG recurrence as well. Black line denotes the initial response. **I,** Cancellation speed as a function of the selectivity of sensory input to dendritic (broad) versus axonal (narrow) spikes, where 1 indicates that sensory input solely effects dendritic spikes and 0 indicates equal effects on broad and narrow spikes. MG cells hinder cancellation when selectivity is low compared to having no MG input at all, (dashed line). This is because if narrow spikes (which are the output signals of the MG cells) are affected by the sensory input, they amplify the response to the EOD in the Output cells. For the colored lines we used the default value of MG->Output coupling. **J**, Cancellation speed as a function of the selectivity of recurrent MG input to dendritic (broad) versus axonal (narrow) spikes. As we show in Figure 6, MG-recurrence speeds cancellation, but this effect requires that recurrent input is non-selective, affecting both narrow and broad spikes. Selectivity = 1 is worse than having no recurrence (dashed line, for the colored lines we used the default value of MG recurrence) because it reduces the magnitude of the negative image transmitted from MG to Output cells.

**Extended data Figure 6.**
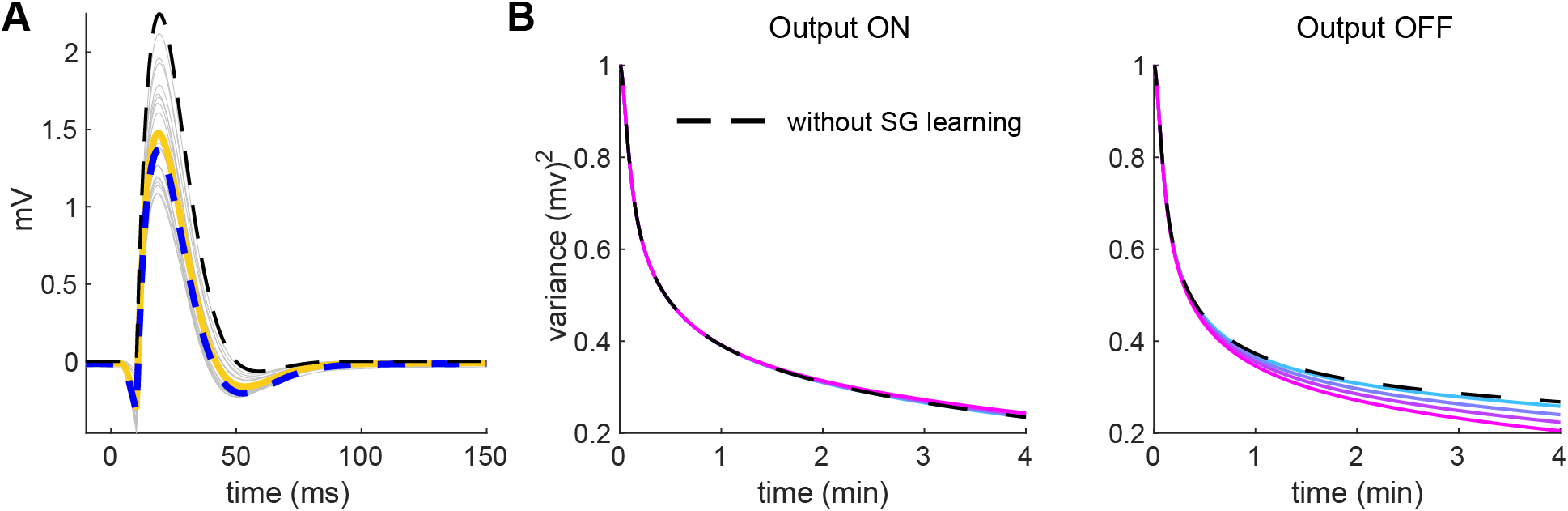
SG cell contribution to sensory cancellation (related to. Figure 6**). A,** Modeling cancellation of predictable sensory input in 20 SG+ cells (gray lines) assuming a uniform distribution of GCA input centered around the median number of GCA-SG synapses found in the connectome (Figure 2). After 20 minutes of learning the 20 SG cells are partially cancelled, and the average value (yellow line) is similar to having one SG cells receiving the average number of GCA inputs (dashed blue line). The dashed black line shows the initial response to sensory input. **B,** Adding learning in SG- does not speed cancellation in Output ON cells, but learning is SG+ cells can speed cancellation in Output OFF cells depending on the strength of their response to sensory input. Colored lines indicate different strengths of the initial response of SG cells to sensory input, with magenta equivalent to measured the MG+ cell response and cyan equivalent to the measured Output ON cell response.

**Extended data Figure 7.**
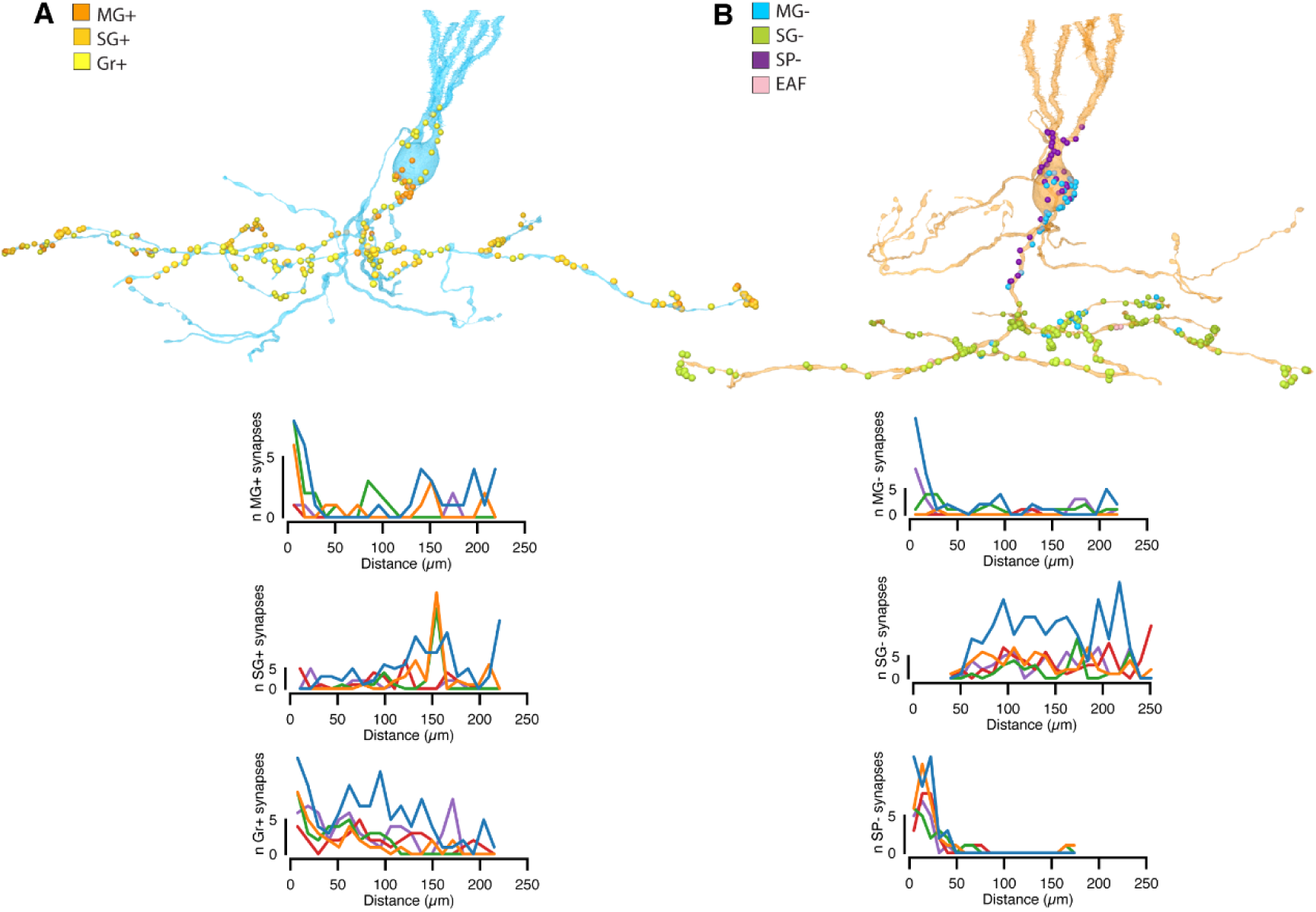
Synaptic placement onto MG cells may support biophysical requirements related to generating and transmitting predictions (related to. Figure 6**). A,** Top, placement of SG+, Gr+, and MG+ synapses (colored circles) onto an MG- cell. Bottom, Distribution of synapse distance from the soma for 5 MG- cells (colored lines). **B**, Top, placement of SG-, Sp-, and MG- synapses (colored circles) onto an MG+ cell. Bottom, Distribution of synapse distance from the soma for 5 M+ cells (colored lines).

## Notes

### Competing Interest Statement

The authors have declared no competing interest.

